# Perivascular tenascin C triggers sequential activation of macrophages and endothelial cells to generate a pro-metastatic vascular niche in the lungs

**DOI:** 10.1101/2021.03.30.436797

**Authors:** Tsunaki Hongu, Maren Pein, Jacob Insua-Rodriguez, Jasmin Meier, Kristin Decker, Arnaud Descot, Angela Riedel, Thordur Oskarsson

## Abstract

When cancers progress to metastasis, disseminated cancer cells frequently lodge near vasculature in secondary organs. However, our understanding of the cellular crosstalk evoked at perivascular sites is still rudimentary. In this study, we identify intercellular machinery governing formation of a pro-metastatic vascular niche during breast cancer colonization in lungs. We show that four secreted factors, INHBB, SCGB3A1, OPG and LAMA1, induced in metastasis-associated endothelial cells (ECs), are essential components of the vascular niche and promote metastasis in mice by enhancing stem cell properties and survival ability of cancer cells. Notably, blocking VEGF, a key regulator of EC behavior, dramatically suppressed EC proliferation, whereas no impact was observed on the expression of the four vascular niche factors in lung ECs. However, perivascular macrophages, activated via the TNC-TLR4 axis, were shown to be crucial for EC-mediated production of niche components. Together, our findings provide mechanistic insights into the formation of vascular niches in metastasis.

## INTRODUCTION

Cancer cell dissemination and development of metastasis in secondary organs is the primary cause of morbidity and mortality in cancer patients. Interactions between disseminated cancer cells and non-transformed stromal cells of secondary organs play a crucial role in metastatic colonization^1^. Numerous stromal cells have been shown to interact with cancer cells leading to generation of a metastatic niche that promotes malignant growth^2, 3^.

Formation of new blood vessels is recognized as a key event to promote cancer growth by delivering nutrition and oxygen to tumors^4^. Vascular endothelial growth factor (VEGF) is a crucial promoter of neoangiogenesis via the VEGF receptor 2 (VEGFR2). VEGF promotes the process by inducing endothelial cell proliferation, migration and survival^5^. This can lead to increased vascular permeability and sprouting by activating tip cells of established vessels^6^. Anti-angiogenic therapy targeting VEGF signaling has been used as a part of therapeutic regimen to treat patients with advanced breast cancer. However, in spite of the importance of VEGF in angiogenesis, the clinical benefits obtained using VEGF neutralization in breast cancer patients have been modest^7^. Importantly, recent findings suggest that blood vessels play a significant role during cancer and metastatic progression that extends beyond delivering nutrients and other essentials to the tumor^8^. Growing evidence suggests that disseminated cancer cells associate with vasculature at the metastatic site, and that enhanced adhesion and crosstalk with endothelial cells (ECs) may regulate phenotype and function of cancer cells in metastasis^9–11^. Whereas, a functional role for endothelium is of critical importance for metastatic progression, our understanding of the complex interactions that occur is still rudimentary.

In this study, we analyzed the interactions between breast cancer cells and ECs in lungs during metastatic progression. We identified four components of a pro-metastatic vascular niche, inhibin beta B (INHBB), secretoglobin family 3A member 1 (SCGB3A1), osteoprotegerin (OPG) and laminin subunit alpha 1 (LAMA1), whose expression associated with poor clinical outcome in breast cancer patients. We show that expression of all four vascular niche components individually promotes metastasis in mice with INHBB and OPG supporting stemness and survival of disseminated cancer cells respectively. Notably, inhibition of VEGF or VEGFR2 did not affect the expression of these four genes in ECs, indicating that the pro-metastatic vascular niche can form independently of VEGF signaling. Exploring the regulatory mechanisms of the vascular niche, we found that it was not directly induced by cancer cells, but via metastasis-associated macrophages as intermediates. Our results indicate that perivascular macrophages, activated by the extracellular matrix protein tenascin C (TNC) via toll-like receptor 4 (TLR4), promote formation of a vascular niche. Importantly, TNC repression, macrophage depletion or TLR4 inhibition in vivo, all suppressed the four niche genes in lung ECs, resulting in disruption of the vascular niche and reduced metastasis. Together, the results reveal a crucial crosstalk within the vascular niche and underscore the importance of extracellular matrix proteins as regulators of the microenvironment in metastasis. The findings suggest that targeting the extracellular matrix or macrophages may be an attractive option to block vascular niche formation and lung metastasis.

## RESULTS

### Endothelial cells undergo marked changes during lung metastasis that are associated with poor outcome in breast cancer patients

To investigate the dynamic molecular changes in ECs during metastatic colonization, we isolated ECs from lungs of mice that had been intravenously injected with MDA231-LM2 breast cancer cells, a highly metastatic derivative of MDA-MB-231 (MDA231) cells^12^. ECs were purified from lungs with growing metastases, at different stages, for transcriptomic studies (Supplementary Figure 1a,b). First, we analyzed expression of endothelial marker CD31 in metastatic nodules at week 1, week 2 or week 3 post cancer cell-injection. This revealed that although early nodules (weeks 1 or 2) grew in proximity of blood vessels, these vessels were most prominent inside the metastatic nodules at later stage (week 3) of metastatic colonization (Figure 1a). CD31-expressing ECs were observed in more than 80% of metastatic nodules at week 3 and this was associated with increased nodule size (Figure 1b,c). The number of ECs was also linked to the number of cancer cells in the lungs, suggesting EC proliferation in addition to vascular cooption in growing metastases (Figure 1d).

**Figure 1.**
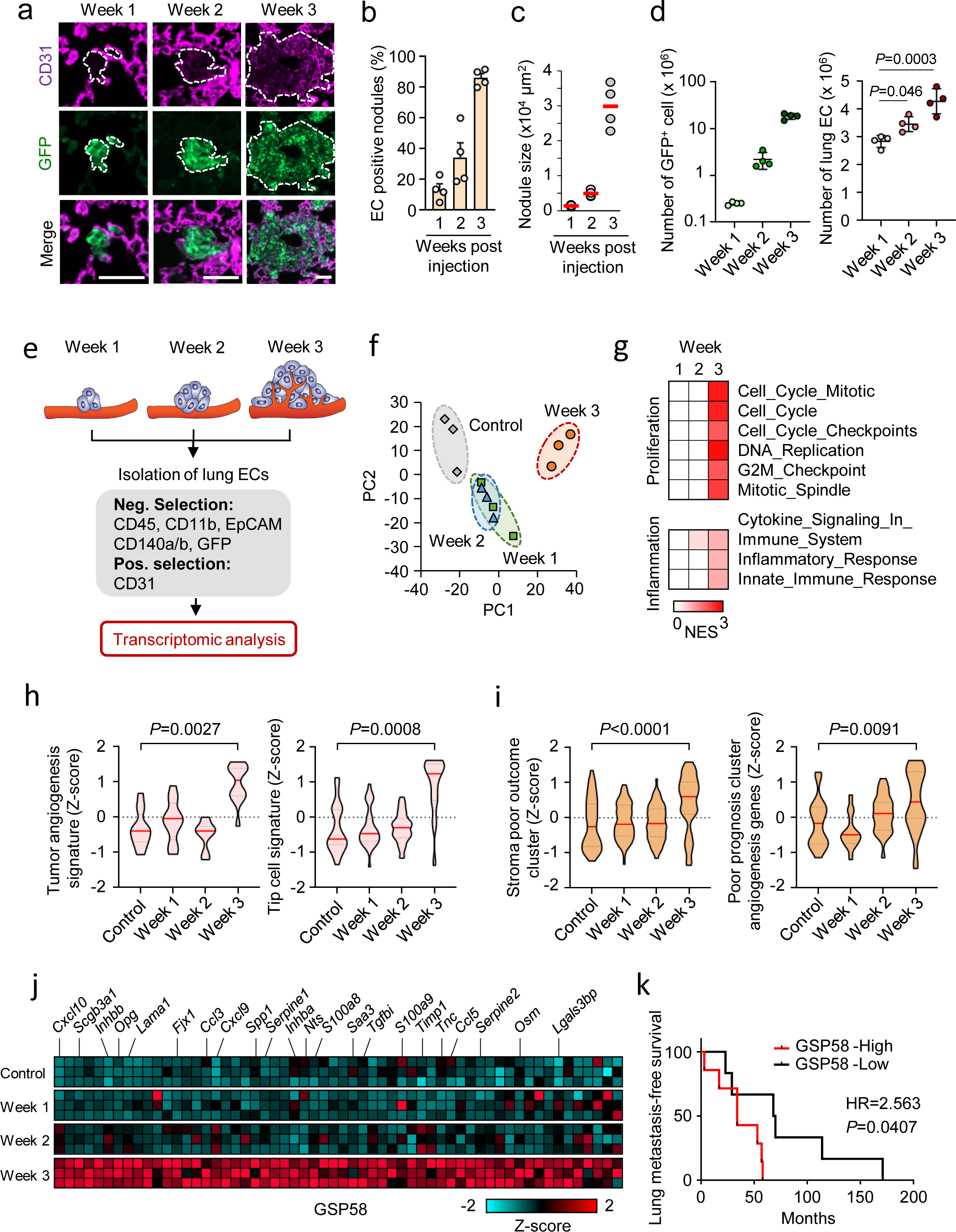
Transcriptomic analysis identifies characteristic changes in reactive ECs during metastatic colonization of the lungs. **a**, Immunofluorescence images showing the association of lung endothelial cells (ECs, CD31) with metastatic breast cancer cells (GFP) in mouse lungs at the indicated time points post intravenous injection of MDA231-LM2 breast cancer cells. Scale bars, 50 μm. Dashed lines show the margins of metastatic foci. **b**, Quantification of metastatic nodules from A that have intra-nodular ECs; *n* = 72 nodules (week 1), 76 nodules (week 2) and 83 nodules (week 3), from 4 mice were analyzed for each time point. Data are mean with SEM. **c**, Size of MDA231-LM2 derived metastatic nodules in lung at week 1-3. Minimum 16 nodules were analyzed for each lung; *n* = 4 mice per each group. **d**, Total number of MDA231-LM2 cancer cells (left) and ECs (right) in lungs at different stages of metastatic colonization. Data show means with SD; *n* = 4 mice for each time point. *P* values were determined by one-way ANOVA with Dunnet’s multiple comparison test. **e**, Experimental setup for EC isolation from mouse lungs at different stage of MDA231-LM2 derived metastasis followed by transcriptomic analysis. **f**, Principal component (PC) analysis of gene expression profiles from ECs isolated from healthy lungs (control) or lungs with different stages of metastasis (as in e). **g**, Gene set enrichment analysis (GSEA) of isolated ECs using several proliferation- or inflammation-related signatures for each time point. Signatures with nominal *P* < 0.05 and FDR < 0.25 were considered significant. NES, normalized enrichment score. **h**, Violin plots showing Z-scores of genes of tumor angiogenesis signature^53^ (left), and tip cell signature^54^ (right) calculated from gene expression profiles of lung ECs isolated from healthy control lungs and metastatic lungs at indicated times. *P* values were determined using averaged Z-scores of each signature by unpaired two-tailed t-test; *n* = 3 each group. **i**, Z-scores of genes from a stroma poor outcome gene cluster^55^ (left) and a poor prognosis angiogenesis gene signature^56^ (right) expressed in lung ECs isolated from mice at indicated time points post intravenous injection of MDA231-LM2 cells. **j**, Heatmap showing the expression of 58 genes encoding secreted proteins (GSP58) that are up-regulated in lung ECs at week 3 post cancer cell injection. Cut-off log2FC > 0.75, P < 0.05, FDRq < 0.25. **k**, Kaplan-Meier analysis of lung metastasis-free survival in breast cancer patients. Lung metastasis samples were stratified according to expression of GSP58 (from panel j) in the metastatic nodules; *n* = 13 human metastasis samples. *P* value was determined by log-rank test. HR, hazard ratio.

To purify ECs from lungs with metastases, we performed flow cytometry to exclude cells expressing makers of hematopoietic -, epithelial -, fibroblastic -, and cancer cells (CD45, CD11b, EpCAM, CD140a/b and GFP) and to select cells expressing the endothelial marker CD31 (Figure 1e, Supplementary Figure 1c). The purity of selected populations was determined by expression of various endothelial markers or distinct markers of other stromal cells (Supplementary Figure 1d,e). Isolated ECs were subjected to transcriptomic analysis using microarrays. Principal component analysis revealed a notable pattern of gene expression changes at different time points of metastasis. Certain changes occur at weeks 1 and 2, but marked differences were not observed between the two time points (Figure 1f). However, the most striking changes occurred in ECs at week 3 that distinguished this time point from the others (Figure 1f). Further analyses, such as gene set enrichment analysis (GSEA) or gene ontology (GO) term analysis, revealed significant changes at week 3 in gene sets regulating cell proliferation or inflammation (Figure 1g; Supplementary Figure 2a,b; Supplementary table 1). Similar results were obtained from Z-score analysis using proliferation- or inflammation-related signatures (Supplementary Figure 2c). Specific targets of transcription factors or pathways involved in these processes such as E2F or Myc (proliferation) or TNF signaling (inflammation) were also prominently upregulated at week 3 (Supplementary Figure 2d). Consistent with these results, angiogenic activity of ECs was significantly promoted at this time point (Figure 1h). Importantly, gene signatures, associated with poor clinical outcome, were induced at week 3, suggesting that endothelial activation is linked to metastatic progression (Figure 1i). Focusing on genes of proteins which can mediate inter-cellular communications, we analyzed expression of extracellular proteins (GO:0005576) in metastasis-associated ECs. We found significant changes in 58 genes of secreted proteins (GSP58) that were particularly induced in ECs at week 3 (Figure 1j; Supplementary table 2). The GSP58 signature was able to cluster samples of lung metastases from breast cancer patients (Supplementary Figure 2e). Moreover, using Kaplan Meier analysis, we observed that high expression of GSP58 in metastases was associated with poor lung metastasis-free survival of breast cancer patients, suggesting that lung ECs may promote metastasis via some of these secreted factors (Figure 1k).

### Induction of GSP58 in reactive endothelial cells is largely independent of VEGF signaling

VEGF signaling is a key pathway regulating angiogenesis in health and disease and targeting VEGF has been explored as a component of metastatic breast cancer therapy^5, 13^. Considering this, we asked the question of whether the changes in genes of ECs in metastasis were affected by VEGF signaling. To address the question, we treated metastases bearing mice with anti-VEGF antibody (B20.4.1.1)^14^ or isotype IgG, two times a week for 3 weeks and isolated lung ECs for transcriptomic analysis (Figure 2a). Common VEGF target genes were repressed in lung ECs and vascular growth was significantly reduced in metastatic lungs, indicating a robust inhibition in VEGF signaling (Supplementary Figure 3a,b). Notably, genes involved in cellular proliferation and angiogenesis were repressed in lung ECs from anti-VEGF treated mice (Figure 2b-e; Supplementary Figure 3c). In addition to gene signatures linked to cell proliferation, such as E2F and Myc targets, gene sets of VEGF signaling targets or VEGF pathway components were among the most repressed gene sets (Figure 2f; Supplementary Figure 3d,e). However, despite the notable repression of VEGF signaling and EC proliferation, gene clusters linked to poor patient outcome were not affected by anti-VEGF treatment (Figure 2g). Importantly, the inflammatory signature and GSP58 were also generally unaffected by the anti-VEGF (Figure 2h,i). Similar results were observed when we analyzed lung ECs from mice treated with anti-VEGF receptor 2 (anti-VEGFR2, DC101) (Supplementary Figure 4a). Expression of selected VEGF target genes in lung ECs was repressed by anti-VEGFR2 treatment (Supplementary Figure 4b). Furthermore, proliferation and angiogenesis signatures were dependent on VEGFR2 (Supplementary Figure 4c-f), but in line with the results from VEGF inhibition, poor outcome gene clusters or GSP58 were not significantly affected by VEGFR2 inhibition (Supplementary Figure 4g,h). These results indicate that whereas anti-VEGF treatment effectively suppresses vascularity within metastatic nodules, the treatment does not repress expression of most of the 58 factors secreted by the vasculature and which associate with poor outcome in patients.

**Figure 2.**
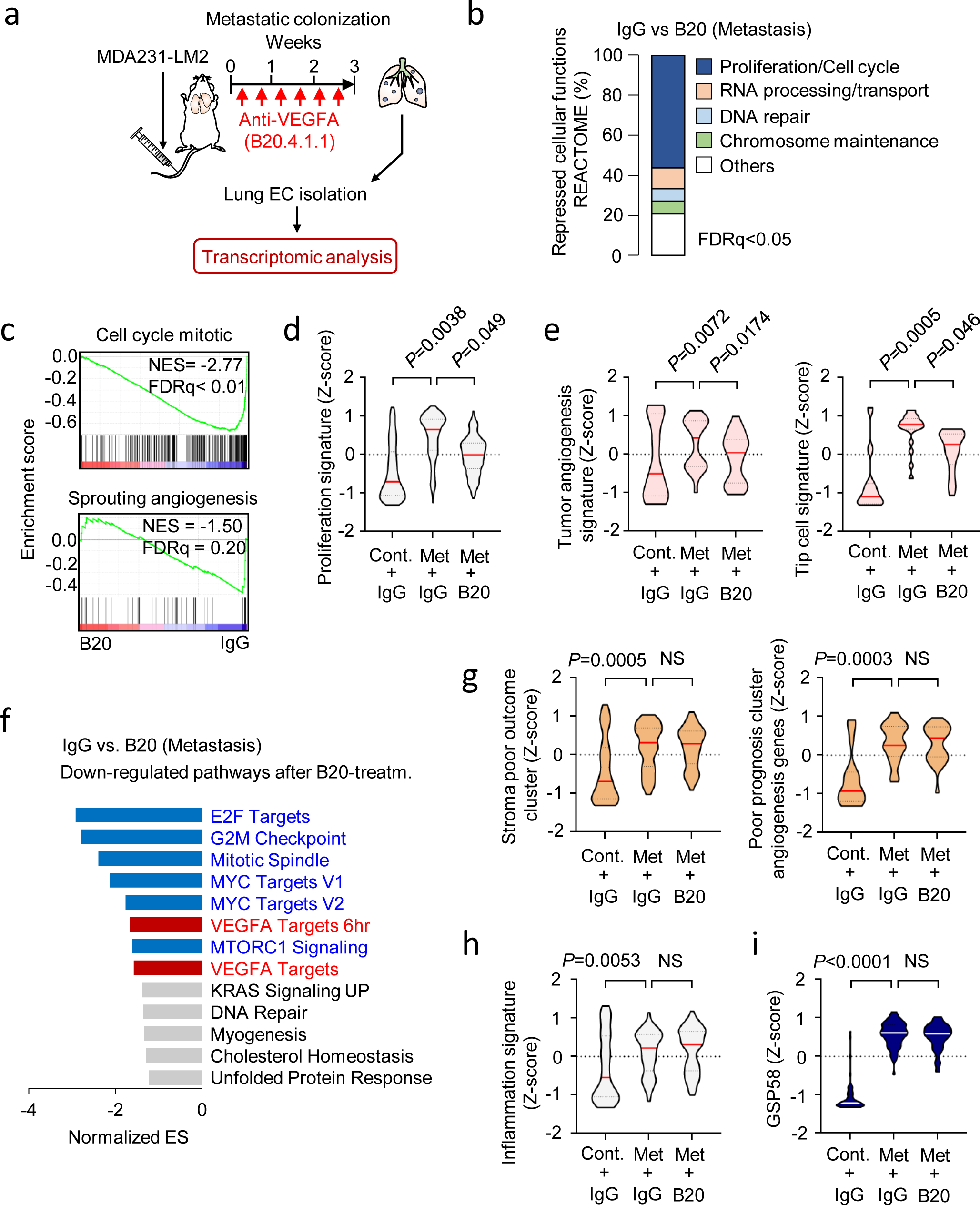
Anti-VEGF treatment inhibits proliferation, but not inflammatory responses or induction of the GSP58 signature in metastasis-associated ECs. **a**, Experimental outline of anti-VEGF treatment of mice with metastatic nodules in the lungs. MDA231-LM2 cells were injected intravenously into NSG mice followed by repeated treatment with anti-VEGF antibody, B20.4.1.1 (B20), or control IgG for 3 weeks. Lung ECs were isolated at week 3 and transcriptomic analysis was performed. Mice harboring comparable lung metastatic loads, measured by *in vivo* bioluminescence imaging, were selected for analysis. **b**, Overview of cellular functions (REACTOME) that are repressed in metastasis-associated ECs by anti-VEGF treatment. Pathways with FDR < 0.05 were included in calculations of percentages. **c**, GSEA of gene signatures, “Cell cycle mitotic” (REACTOME) and “Sprouting angiogenesis” (C5 collection in MSigDB), in lung ECs after B20-treatment compared to control IgG. NES, normalized enrichment score. FDR was determined from *P* values calculated by random permutation test. **d-e**, Violin plots showing Z-scores of genes from a proliferation signature (Cell cycle mitotic, REACTOME) (d), tumor angiogenesis signature^57^ (e, left) and tip cell signature^54^ (e, right) expressed in purified lung ECs from control and metastasis bearing mice with indicated treatment. **f**, Downregulated gene signatures in metastasis-associated ECs treated with B20 compared to IgG control, based on GSEA. Signatures with FDR < 0.25 are shown. Proliferation-related signatures, blue; signatures of VEGF target genes, red; others, grey. **g-i**, Violin plot analysis of stroma poor outcome cluster^55^ (g, left) and angiogenesis genes associated with poor prognosis ^56^ (g, right), inflammation signature (Inflammatory response, Hallmark in MSigDB) (h) and GSP58 (i) expressed in ECs from lungs of mice under indicated conditions. *P* values in d, e and g-i were determined with averaged Z-score of genes in signatures by one-way ANOVA with Fisher’s LSD test; *n* = 3 mice for each group. NS, not significant.

### Inhbb, Lama1, Scgb3a1 and Opg produced by reactive ECs promote lung metastasis

To identify putative gene candidates from GSP58 that are particularly relevant for ECs as a cellular source compared to other stromal cells, we investigated the most highly induced genes within the signature and analyzed their expression in four different stromal cell types isolated from mouse lungs harboring growing metastases (Figure 3a; Supplementary Figure 5a). Expression analysis in lung ECs, fibroblasts, hematopoietic cells and epithelial cells revealed four genes within GSP58, *Inhbb*, *Lama1*, *Scgb3a1* and *Opg*, that were distinctly expressed in ECs compared to other cell types (Figure 3a; Supplementary Figure 5b). Moreover, we confirmed that these four genes were not significantly expressed in cancer cells, but particularly induced in lung ECs at week 3 and unaffected by anti-VEGF treatment (Supplementary Figure 5c-e). We revealed consistent induction of *Inhbb*, *Scgb3a1*, *Opg* and *Lama1* in lung ECs from two xenograft models of lung metastases (MDA231-LM2/SUM159-LM1) and two syngeneic mouse models (4T1/E0771) (Figure 3b). Notably, expression of the four candidate genes correlated significantly with expression of the endothelial maker *Cdh5* (VE-cadherin) in samples of human breast cancer metastases (Figure 3c). This encouraged us to study the function of the four candidate genes in metastasis. Thus, to address whether *Inhbb*, *Scgb3a1*, *Opg* and *Lama1* could induce metastasis in lungs, we ectopically expressed the genes individually in human breast cancer cells and injected intravenously into NSG mice. We expressed the genes in SUM159 and MDA-MB-231 (MDA231), the parental cells of SUM159-LM1 and MDA231-LM2 respectively (Supplementary Figure 5f). Expression of all four candidates individually promoted metastatic colonization of the lungs by SUM159 and MDA231 breast cancer cells (Figure 3d-f). Moreover, combined expression of the four genes in MDA231 cancer cells showed an additive induction of metastatic growth in lungs (Figure 3e,f). This suggested that as individual genes, *Inhbb*, *Scgb3a1*, *Opg* and *Lama1* can indeed promote metastatic colonization. Moreover, coexpression of the four genes may provide a further advantage to the cancer cells. The results indicate that proteins encoded by the four gene candidates are functional components of a pro-metastatic vascular niche. To address the potential association with clinical outcome, we performed Kaplan Meier analyses using expression of the four vascular niche components in estrogen receptor (ER)-negative breast cancer samples and investigated the potential link to survival. In these samples, expression of the four vascular niche factors was significantly associated with poor relapse-free and overall survival, indicating a potential role in breast cancer patients (Figure 3g).

**Figure 3.**
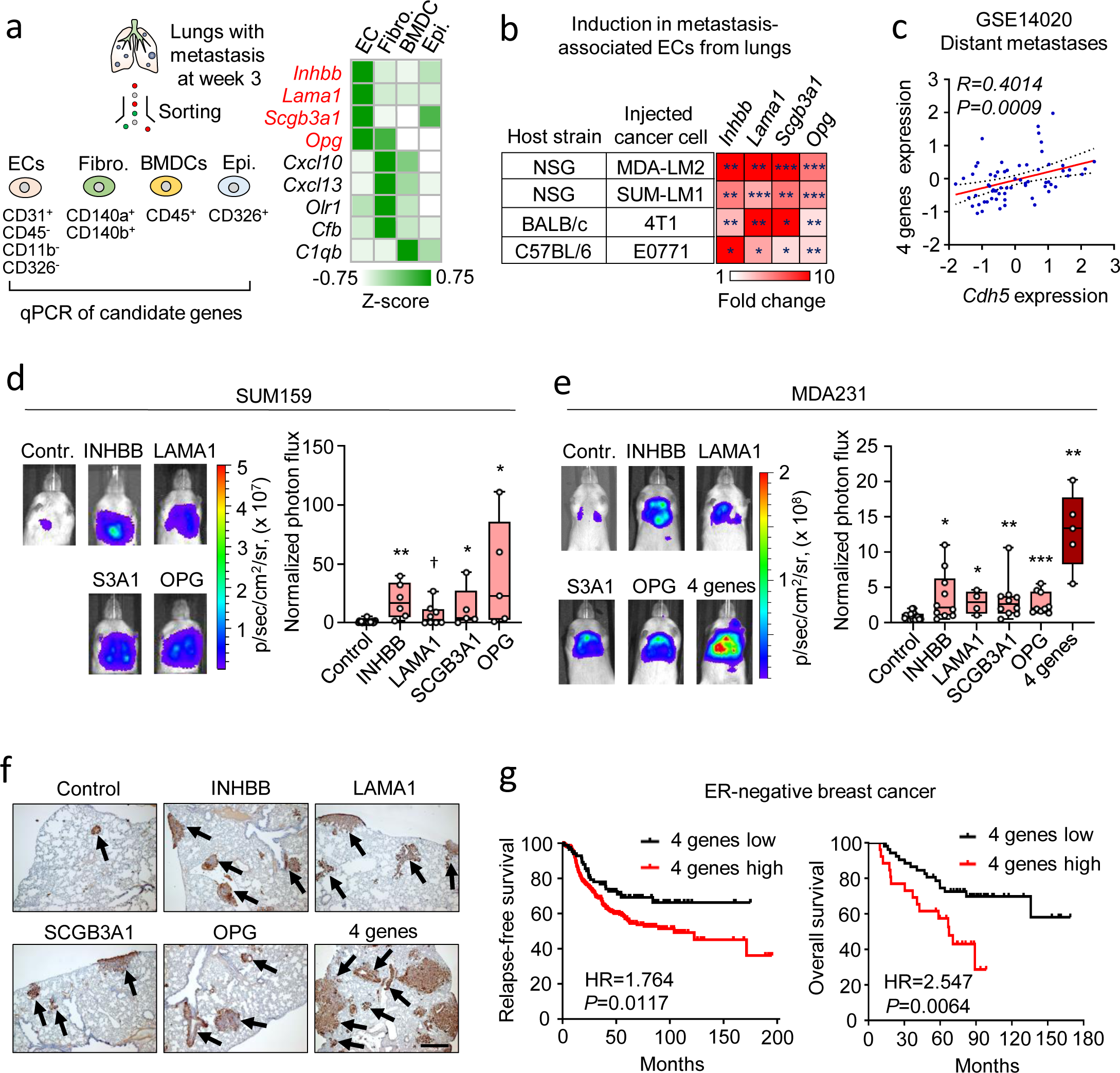
Secreted factors produced by reactive endothelial cells promote lung metastasis. **a**, Schematic of the analysis of endothelial niche candidate genes in different stromal cells isolated from metastatic lungs (at week 3, left) and heatmap summarizing the expression of the top genes with log2FC >3.5 (right). Gene expression is shown as a Z-score calculated from the results of qPCR of stromal cells isolated from four mice per group. The four genes highly expressed in ECs are highlighted in red. **b**, Expression of four endothelial niche candidate genes in lung ECs isolated from indicated mouse strains with metastasis of different breast cancer cell lines. Heatmap was generated from qPCR analysis of the four genes comparing with healthy lung from a corresponding strain. *P* values were calculated by unpaired one-tailed t-test. * *P* < 0.05, ** *P* < 0.01 and *** *P* < 0.001; *n* = 3-4 mice per group. **c**, Correlation analysis of normalized expression of the four candidate genes and *Cdh5* in metastatic lesions of breast cancer patients (GSE14020). Linear regression with Pearson correlation r and two-tailed *P* value is shown. *n* = 65. **d-e**, Metastatic colonization of the lungs by SUM159 breast cancer cells (d) and MDA231 cells (e) overexpressing endothelial niche factors or a control vector. Bioluminescence images (left) and normalized photon flux (right) 42 days post intravenous injection are shown. Boxes depict median with upper and lower quartiles. The data points show values of biological replicates and whiskers indicate minimum and maximum values. *P* values were calculated by one-tailed Mann-Whitney test between overexpression and empty control. * *P* < 0.05, ** *P* < 0.01, and *** *P* < 0.001 for d and e. ^†^ *P* = 0.069. **f**, Histological examples of metastases marked by expression of human vimentin in lungs of mice injected with MDA231 cells overexpressing endothelial niche factors or a control vector. Scale bar, 200 μm. **g**, Kaplan-Meier analyses of ER-negative breast cancer patients with KM plotter, associating expression of four endothelial niche factors with relapse-free survival (left, *n* = 347 patients) and overall survival (right, n = 79 patients). *P* values were determined by log-rank test. HR, hazard ratio.

### INHBB promotes stem cell properties and OPG protects breast cancer cells from TRAIL-induced apoptosis

We explored the cellular function of two of the vascular niche factors to understand their role in metastasis. To determine the function of INHBB, we performed transcriptomic analysis on cancer cells treated with activin B, a growth factor that is composed of an INHBB homodimer (Figure 4a). This revealed a response gene signature that associated with poor relapse-free survival in breast cancer patients (Figure 4b,c; Supplementary table 3). Within the signature, three members of the family of inhibitors of DNA binding and cell differentiation (ID) proteins (ID1, ID2 and ID3) were among the top induced genes (Figure 4b). These proteins are recognized to promote stem cell self-renewal and multipotency and were shown to facilitate metastatic colonization by breast cancer cells^15, 16^. To analyze the phenotype of activin B-stimulated cancer cells, we stratified gene expression datasets from samples of breast cancer patients (Metabric or samples of lung metastases) based on expression levels of the activin B response signature (Figure 4d). Then we performed GSEA on activin B signature high or low patient samples to address whether the two groups had distinct properties. We observed that patients with high expression of activin B signature exhibited enrichment in a number of stem cell signatures (Figure 4e). Following up on these results, we analyzed sphere formation under ultra-low adhesion conditions, a recognized assay that can determine and enrich for stem cells of many tissues^17, 18^. First, we observed that conditioned medium from cultured lung ECs can stimulate oncosphere formation by two metastatic breast cancer cell lines, SUM159-LM1 and MDA231-LM2 (Figure 4f). Moreover, recombinant activin B significantly stimulated oncosphere formation in breast cancer cells (Figure 4g). Finally, conditioned medium obtained from lung ECs transduced with shRNA against *Inhbb*, showed reduced ability to stimulate oncosphere formation compared to control conditioned medium (Figure 4h). Together, these experiments suggest an important role for INHBB in promoting stem cell properties in breast cancer cells.

**Figure 4.**
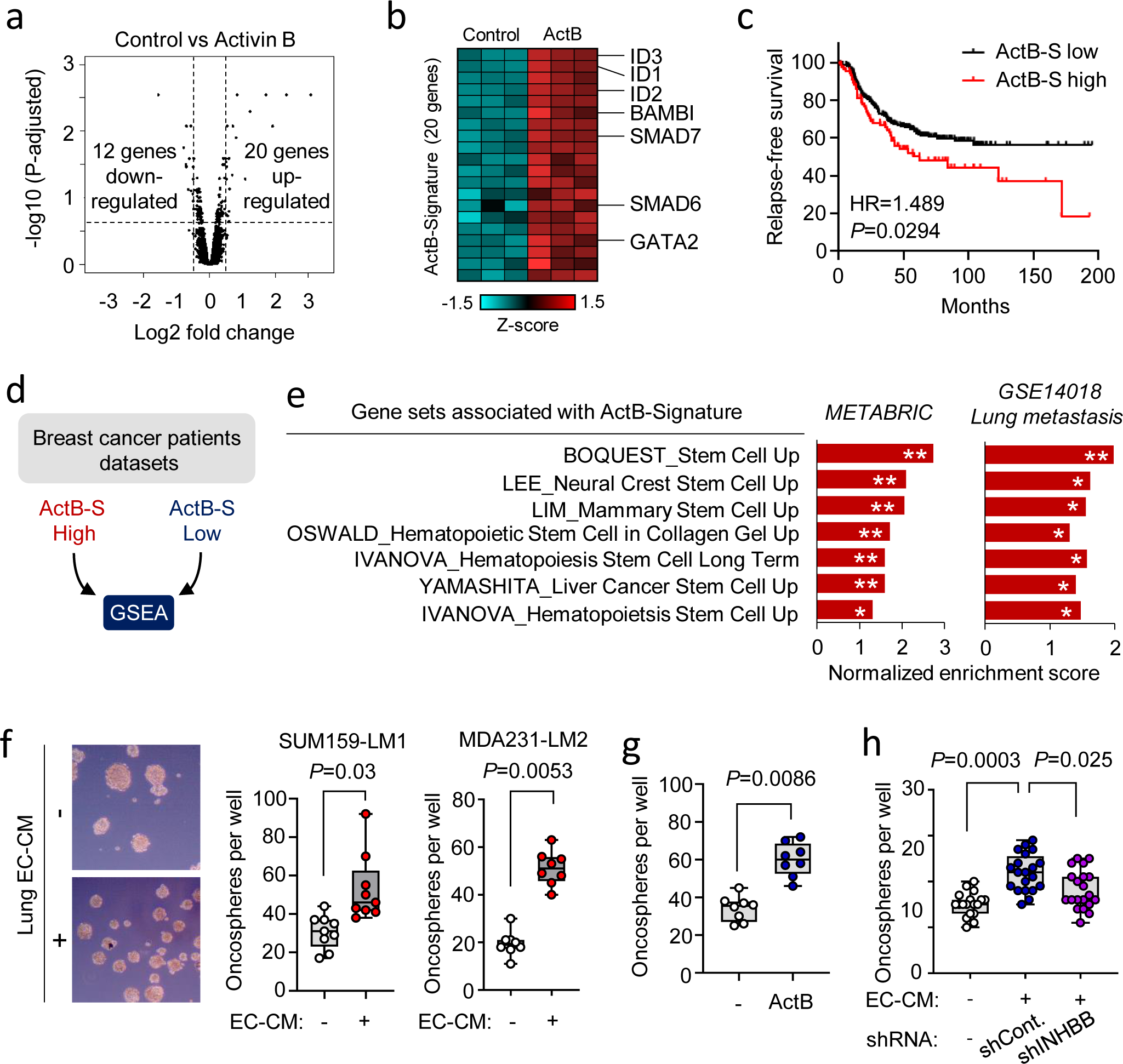
The INHBB homodimer, activin B, promotes stem cell properties of breast cancer cells. **a**, Volcano plot showing differential gene expression in SUM159-LM1 breast cancer cells treated with 50 ng/ml of the INHBB homodimer, activin B, for 6 h compared to control; *n* = 3 biological replicates. **b**, Heatmap of the top 20 upregulated genes (activin B-signature). **c**, Kaplan-Meier analysis of relapse-free survival of ER-negative breast cancer patients stratified according to expression of activin B-signature (n = 347 patients). *P* value was determined by log-rank test. HR, hazard ratio. **d-e**, GSEA of datasets from breast cancer patients that were divided into activin B-signature high and low groups. Schematic diagram (d) and stem cell-associated gene sets enriched in patients with high activin B-signature expression (e). Dataset from triple negative breast cancer samples in METABRIC discovery (upper and lower quantile; *n* = 46) and lung metastases of breast cancer in GSE14018 (*n* = 16) were analyzed. FDR was determined from *P* values calculated by random permutation test. * FDR < 0.25, ** FDR < 0.05. **f**, Oncosphere formation of SUM159-LM1 (pictures and left graph) and MDA231-LM2 (right graph) breast cancer cells cultured with conditioned medium (CM) from ST1.6R human lung ECs. *P* values were calculated by ratio-paired two-tailed t-test with 3 independent experiments. **g**, Oncosphere formation of SUM159-LM1 stimulated with 50 ng/ml of recombinant activin B. *P* value was determined by ratio-paired two-tailed t-test with 4 independent experiments. **h**, Oncosphere formation of SUM159-LM1 cultured with CM derived from ST1.6R transduced with shControl or shINHBB. Data shown are from 5 independent experiments and *P* values were determined by one-way ANOVA with Dunnet’s multiple comparison test. Each dot in f-h indicates all technical replicates of independent experiments, boxes depict median with upper and lower quartiles, whiskers indicate minimum and maximum values.

OPG is a protein that can function as a decoy receptor for TRAIL and neutralize its function^19^. TRAIL is highly expressed in the lung microenvironment (Figure 5a) and has been recognized to be an important regulator of apoptosis in the lung during metastasis^20^. We analyzed TRAIL-induced apoptosis of the MDA231-LM2 cells by cleaved caspase 3 protein expression in the presence of incremental levels of OPG, and observed a significant OPG mediated protection from TRAIL and thus reduced apoptosis (Figure 5b). This suggested that OPG expression induced in metastasis-associated ECs may protect the invading cancer cells from TRAIL-induced apoptosis. Indeed, in mice we observed that metastases expressing OPG exhibited less apoptosis compared to control, based on cleaved caspase 3 expression (Figure 5c). The receptors that bind and mediate TRAIL-induced apoptosis are the death receptors 4 and 5 (DR4/5)^21, 22^. To address the significance of these receptors for metastatic colonization of the lungs, we induced shRNA-mediated knockdown of DR4/5 in MDA231 breast cancer cells (Supplementary Figure 5g) and injected intravenously into NSG mice. DR4/5 knockdown significantly promoted metastatic colonization of the lung (Figure 5d) and exhibited less apoptosis in the lung nodules (Figure 5e), suggesting that this mechanism may play an important role in protecting the lung against metastatic colonization. Taken together, the results on INHBB and OPG function suggest that they respectively promote stem cell properties and survival in response to TRAIL.

**Figure 5.**
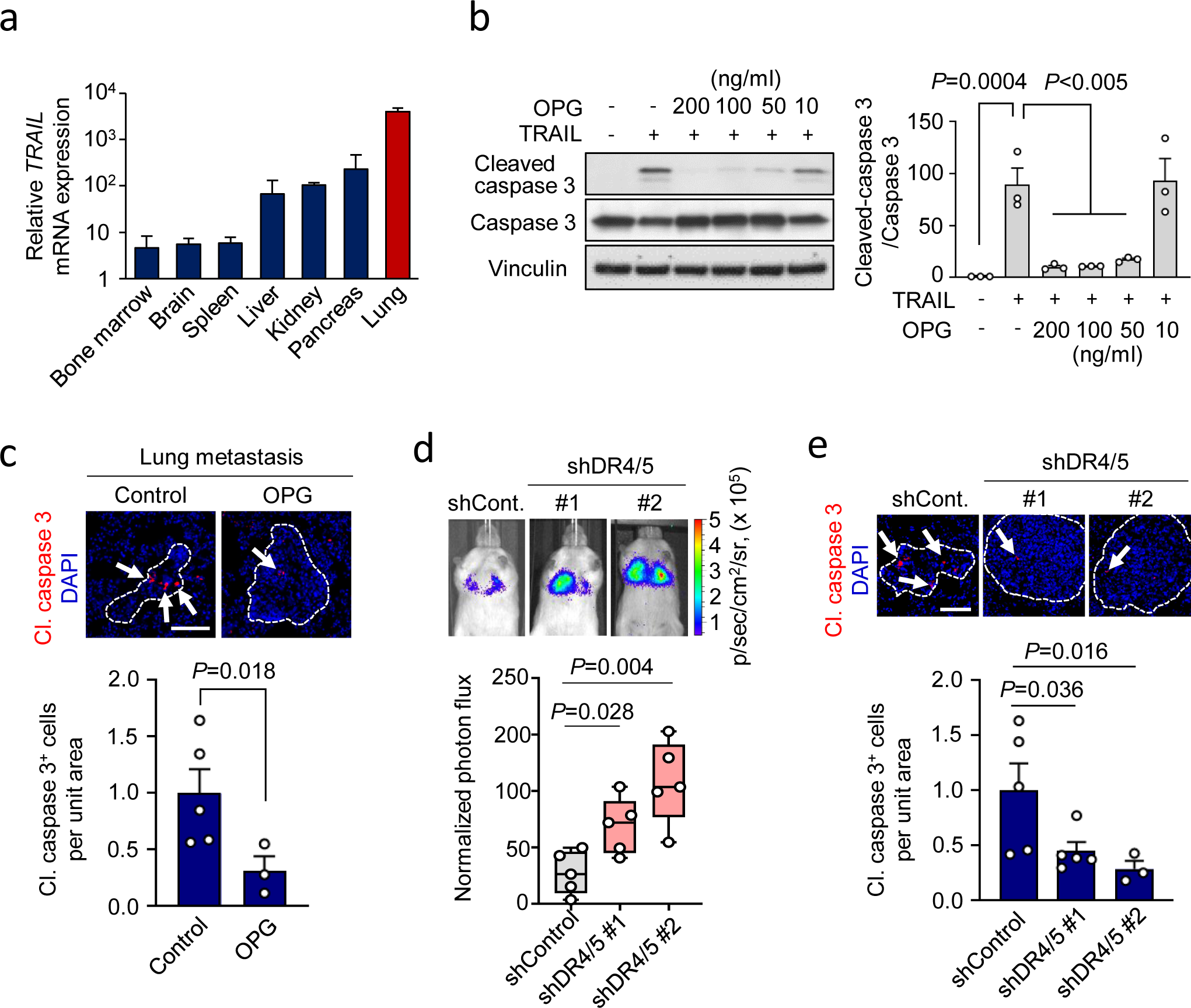
OPG regulates breast cancer cell survival. **a**, TRAIL mRNA levels in different organ tissues isolated from NSG mice; *n* = 3 mice per group. Expression was determined by qPCR. **b**, Left, Western blot analysis of cleaved caspase 3 expression in MDA231-LM2 cells treated with TRAIL (50 ng/ml) alone or in combination with indicated concentrations of OPG. Right, quantification based on the ratio between cleaved caspase 3 and uncleaved caspase 3. Means with SEM calculated from 3 independent experiments are shown. Statistical analysis was performed with one-way ANOVA with Dunnet’s multiple comparison test. **c**, Immunofluorescence analysis of cleaved caspase 3 expression in lung metastasis from mice injected with control or OPG transduced MDA231 breast cancer cells. Top, representative examples; cleaved caspase 3 (red), DAPI for nuclear staining (blue). Scale bar, 100 μm. Bottom, quantification; *n* = 5 for control and 3 for OPG. *P* value was determined by one-tailed Mann-Whitney test. **d**, Lung colonization of DR4 and DR5 knockdown MDA231 cells. Bioluminescence was used to determine the metastatic burden in the lungs. Representative bioluminescence images (top) and normalized photon flux (bottom) 21 days after intravenous injection are shown; *n* = 5 mice per group. Boxes show median with upper and lower quartiles and whiskers indicate maximum and minimum. *P* values were calculated by one-tailed Mann-Whitney test. **e**, Cleaved caspase 3 analysis of lung metastasis with control or DR4/5 shRNAs. Top, representative examples; cleaved caspase 3 (red), DAPI for nuclear staining (blue). Scale bar, 100 μm. Bottom, quantification; *n* = 3-5 mice per group. *P* values were determined by one-tailed Mann-Whitney test.

### The vascular niche is regulated by macrophages in metastatic lungs

We hypothesized that cancer cells could directly be responsible for the induction of the vascular niche components. Therefore, we treated lung ECs with conditioned medium from MDA231-LM2 breast cancer cells and analyzed expression of niche factors in ECs. Conditioned medium from cancer cells did not promote expression of the vascular niche factors in ECs (Supplementary Figure 6a), indicating that a different cell type was likely responsible for the induction. With this in mind, we used expression of GSP58 to stratify samples of lung metastases from breast cancer patients and analyzed the properties of each group using GSEA (Figure 6a). We observed that metastatic samples that expressed GSP58 exhibited enrichment of gene signatures that are linked to innate immune cells (Figure 6b). Thus, we examined whether these cells might be inducers of the vascular niche and selected neutrophils or macrophages for further analysis. Conditioned medium from an activated macrophage cell line, but not a neutrophil cell line, indeed promoted expression of the niche components in lung ECs (Figure 6c and Supplementary Figure 6b). This suggested that macrophages may serve as intermediate for metastasis induced changes in ECs and formation of a vascular niche. We used immunofluorescence to address the recruitment of F4/80 expressing macrophages to metastatic nodules in lung. Indeed, a substantial number of macrophages infiltrating the metastatic nodules was observed (Supplementary Figure 6c). Notably, all metastatic nodules that we investigated contained macrophages, which were frequently localized to blood vessels (Figure 6d,e). To investigate the functional role of macrophages in endothelial activation and metastatic colonization, we transiently depleted macrophages from mice harboring growing metastases, using clodronate-liposome (Figure 6f). This blunted the increase in macrophages observed in metastatic mouse lungs (Supplementary Figure 6d). We isolated lung ECs from these mice and performed transcriptomic analysis. Importantly, macrophage depletion repressed expression of numerous genes within GSP58 that were induced in ECs from metastatic lungs (Figure 6g,h). The GSP58 changes, in the absence of macrophages, included a significant reduction in the niche genes *Inhbb*, *Lama1*, *Scgb3a1* and *Opg* (Figure 6i). Interestingly, whereas expression of a proliferation gene signature was unaffected by macrophage depletion, the EC inflammatory responses were repressed (Supplementary Figure 6e-h). GSEA revealed that inflammation-related signaling, such as TNF-, IL6- and IL1-mediated pathways, were prominently repressed in ECs from metastatic lungs in which macrophages had been depleted (Supplementary Figure 6i). These results suggest that metastasis-associated macrophages induce inflammatory responses in lung ECs. In line with the results from xenograft mouse models, the elimination of macrophages in the fully immunocompetent 4T1 mouse mammary tumor model also repressed the expression of the four niche genes, indicating that macrophages are a crucial regulator of vascular niche factors in the context of intact immune system (Supplementary Figure 7a-c). Moreover, consistent with the *in vitro* results, no change in niche factor expression was observed following elimination of neutrophils in the 4T1 model (Supplementary Figure 7d-f). Finally, long-term depletion of macrophages significantly repressed metastatic colonization of the lung by MDA231-LM2 cancer cells (Figure 6j). Together, the results indicate that perivascular macrophages function as a key regulator of the vascular niche during lung metastasis.

**Figure 6.**
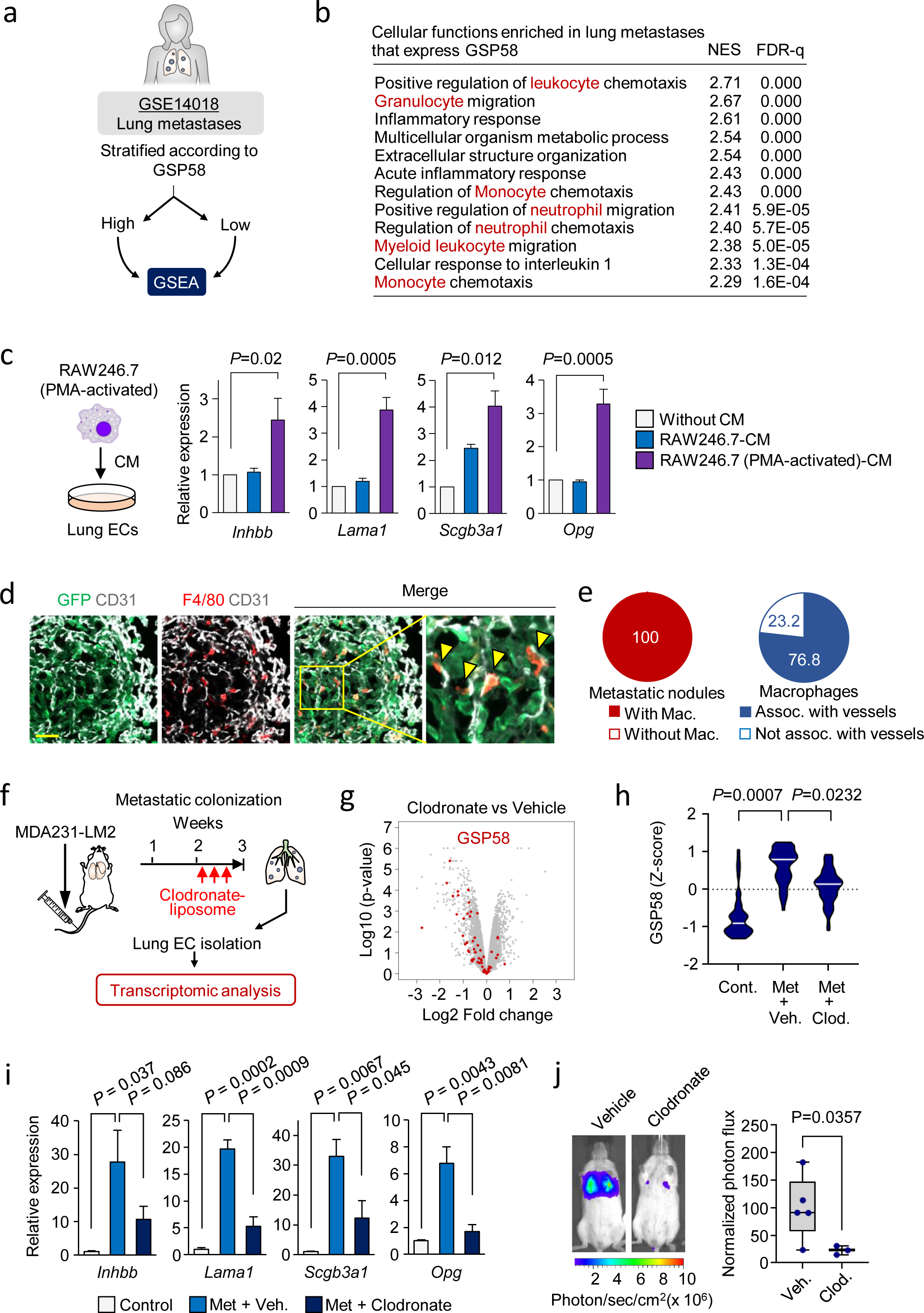
The endothelial niche is regulated by perivascular macrophages in lung metastasis. **a**, Schematic overview of the GSEA setup to analyze datasets from human metastases samples ranked according to GSP58. All patients with lung metastasis (16 patients) were selected from GSE14018. This analysis was performed to identify potential cellular sources of EC activation since cancer cells were not able to do it directly. **b**, Cellular functions enriched in human lung metastases that express GSP58. The cellular functions are ranked based on normalized enrichment scores (NES). FDR was determined from *P* values calculated by random permutation tests. **c**, Expression of 4 factors of the endothelial niche in ECs treated with conditioned media from naïve or activated macrophage cell line RAW246.7. Means with SEM from 4 or 5 independent experiments are shown. Statistical analysis was performed with ratio-paired two-tailed t-tests. **d**,**e**, Immunofluorescence analysis of endothelial cells (CD31, white), macrophages (F4/80, red) and MDA231-LM2 (GFP, green) in metastatic nodules from mouse lungs. Shown are representative images (d). Arrow heads indicate perivascular localization of macrophages in a metastatic nodule. Scale bar, 50 μm. Quantification (percentages) of nodules with infiltrated macrophages or macrophages associated with vessels (e). **f**, Experimental setup of Clodronate-mediated macrophage depletion in mice with metastasis in lungs, followed by transcriptional analysis of lung ECs. Mice harboring comparable lung metastatic loads, measured by *in vivo* bioluminescence imaging, were selected for analysis. **g**, Volcano plot showing differential expression of genes in lung ECs after macrophage depletion. GSP58 are highlighted in red color. **h**, Violin plot showing expression of GSP58 signature in ECs from lungs of metastasis-bearing mice following macrophage depletion. *P* value were determined by one-way ANOVA with Fisher’s LSD test. **i**, Expression of the four endothelial niche factors in purified lung ECs as in F-H. mRNA levels were determined by qPCR; *n* = 3 mice. *P* values were calculated by one-way ANOVA with Dunnet’s multiple comparison test. **j**, Lung colonization of MDA231-LM2 cells in mice after long term treatment with clodronate-liposome. Bioluminescence (left) and photon flux of day 14 normalized to day 7 (right); *n* = 5 mice for PBS-liposome (vehicle), and *n* = 3 mice for clodronate-liposome. Boxes show the median value with upper and lower quartiles. Whiskers represent minimum and maximum values. *P* value was calculated by one-tailed Mann-Whitney test.

### TNC-TLR4 axis promotes activation of perivascular macrophages and the subsequent formation of a pro-metastatic endothelial niche in the lungs

The extracellular matrix is increasingly recognized as an important regulator of cancer progression and metastasis^23–25^. Tenascin C (TNC) is an extracellular matrix protein that has been previously shown to be expressed by breast cancer cells as a crucial component of the metastatic niche in the lung^26^ and (Supplementary Figure 8a). We explored the localization of TNC with respect to metastasis-associated macrophages and endothelium. Immunofluorescence analysis showed that TNC co-localized with vasculature and with perivascular macrophages in metastatic nodules (Figure 7a). Moreover, in human metastases isolated from breast cancer patients, TNC expression or expression of a gene signature from activated macrophages^27^ overlapped with expression of GSP58 and with each other in the patient samples and showed a significant correlation (Figure 7b,c and Supplementary Figure 8b). This suggested a link between TNC expression, metastasis-associated macrophages and activation of a vascular niche. Interestingly, previous studies have shown that TNC can bind and activate toll-like receptor 4 (TLR4) in different cell types, including macrophages, leading to inflammatory signaling in mouse models of arthritis^28^. In line with these findings, we observed that macrophages treated with recombinant TNC-induced expression of *TNF*, *Il1b* and *Il6*, markers of macrophage activation, and this was reverted by inhibition of TLR4 (Figure 7d). However, recombinant TNC did not directly induce expression of the vascular niche proteins in ECs (Supplementary Figure 8c). Considering these results, we hypothesized that perivascular TNC might be involved in activation of macrophages to induce expression of vascular niche components in the lung ECs during metastasis. To address this *in vivo*, we injected control or TNC knockdown MDA231-LM2 breast cancer cells intravenously into NSG mice (Figure 7e, top). Consistent with previous studies, TNC deficiency also reduced metastatic colonization ability of the breast cancer cells (Figure 7f). We isolated ECs from these mice using flow cytometry and determined the expression of the vascular niche genes. All four genes were induced in metastatic foci in a TNC dependent manner (Figure 7g). Moreover, we addressed the role of TLR4 in these processes by treating metastases bearing mice with TLR4 inhibitor (TLR4i) and isolating ECs for analysis (Figure 7e, bottom). Treatment with TLR4i significantly reduced the activation of macrophages, based on TNF-α expression (Supplementary Figure 8d). However, macrophage recruitment to metastatic nodules was unaffected by TLR4i (Supplementary Figure 8e). Importantly, similar to the results in TNC knockdowns, the inhibition of TLR4 caused downregulation of the vascular niche components that were induced in metastatic nodules and impeded metastatic colonization of the lung (Figure 7h left, 7i and Supplementary Figure 8f). Considering the crosstalk between innate and adaptive immunity, we addressed the functional role of TLR4 in a fully immunocompetent mammary tumor model where 4T1 mammary tumor cells are injected into BALBc mice. In line with the results from the xenograft models, TLR4 treatment significantly inhibited metastatic colonization in the 4T1 model (Figure 7h right and Supplementary Figure 8g). The results were crucially reproduced in an orthotopic mouse model of spontaneous metastasis to lung (Figure 7j,k). Whereas mammary tumor growth was not significantly affected by TLR4 inhibition (Supplementary Figure 8h), both size and number of metastatic nodules were reduced upon TLR4i treatment (Figure 7k). Together, these results suggest a microenvironmental interactions in metastasis, where cancer cells produce TNC that activates macrophages which in turn promote the formation of a pro-metastatic vascular niche (Figure 7l).

**Figure 7.**
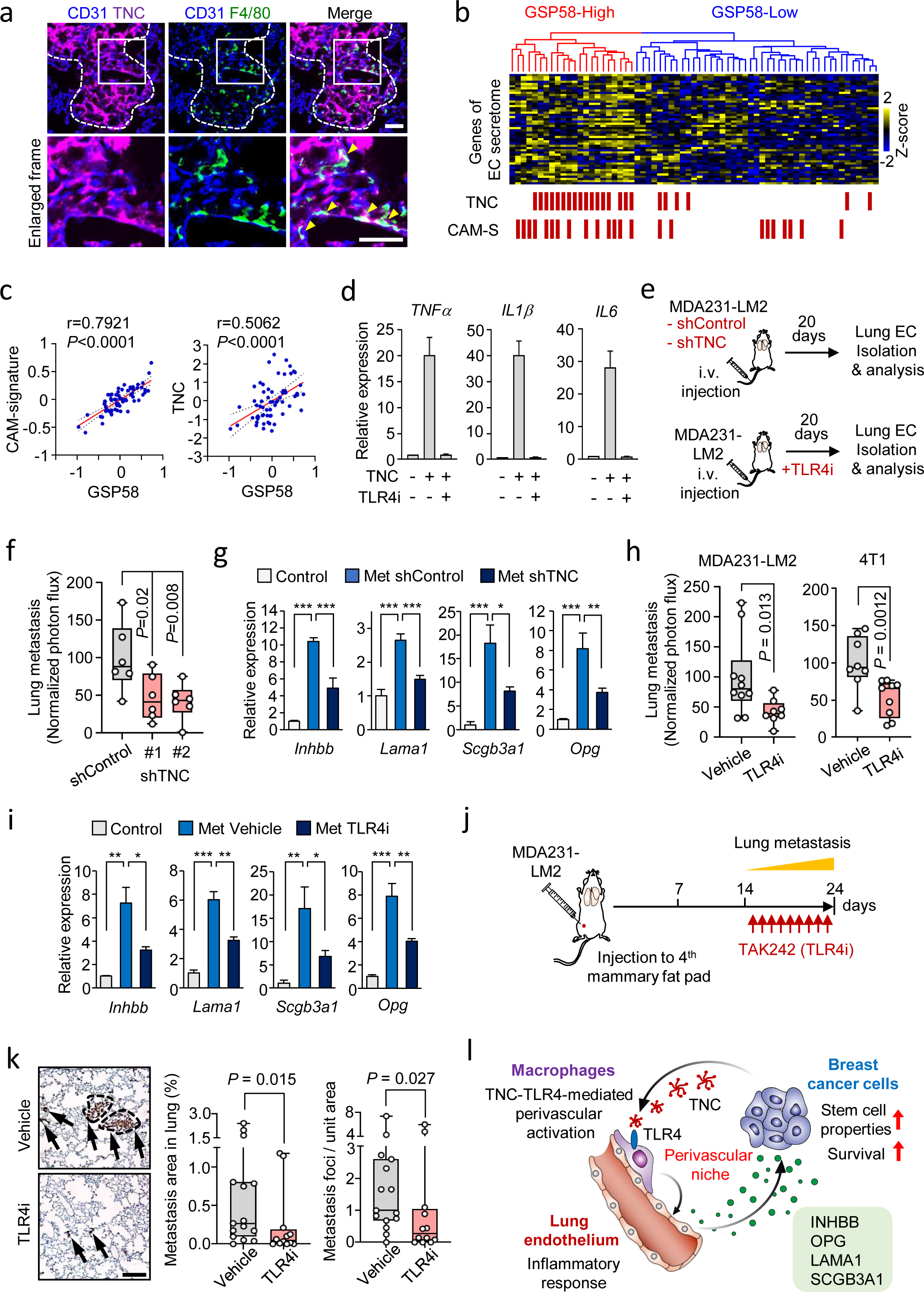
TNC-TLR4 axis promotes activation of perivascular macrophages and the subsequent formation of a pro-metastatic endothelial niche in the lungs. **a**, Immunofluorescence analysis of TNC (purple), macrophages (F4/80, green) and ECs (CD31, blue) in a metastatic nodule from lungs of mice intravenously injected with MDA231-LM2 breast cancer cells. Arrow heads indicate the co-localization of TNC and macrophages at the perivasculature. Dashed line shows the margin of metastatic foci. Scale bar, 50 μm. **b**, Unsupervised hierarchical clustering of 65 human metastases of breast cancer (GSE14020) according to the expression of GSP58. Status of TNC expression and signature of classically activated macrophages (CAM-S) are depicted below the heatmap with red squares, indicating TNC-positive or CAM-S-positive metastasis samples. **c**, Correlation analysis of indicated parameters in 65 metastases samples from breast cancer patients. Linear regression with Pearson correlation r and two-tailed *P* values are shown. **d**, Expression of indicated markers in macrophages isolated from bone marrow and treated with recombinant TNC or the combination of TNC and TLR4 inhibitor (TLR4i, TAK-242) for 6h; *n* = 4 independent experiments **e**, Schematic of experimental outlines where control or shTNC transduced MDA231-LM2 cancer cells were injected intravenously into mice (top) or where mice injected intravenously with MDA231-LM2 cells were treated with TLR4i (bottom). For both experiments, metastatic burden was determined and lung ECs isolated for gene expression analysis. **f**, Lung metastatic colonization quantified based on bioluminescence in control and two independent TNC knockdown breast cancer cells; *n* = 6 mice for each group. *P* values were calculated by one-tailed Mann-Whitney test. **g**, Expression of the 4 factors of the perivascular niche from ECs isolated from metastatic lungs as in f; *n* = 4 mice. *P* values were determined by one-way ANOVA with Dunnet’s multiple comparison test. * *P* < 0.05, ** *P* < 0.01, *** *P* < 0.001. **h**, Bioluminescence analysis of lungs from mice harboring lung metastases and treated with TLR4i. For MDA231-LM2, control: n = 10 mice, TLR4i: n = 8 mice. For 4T1, control: *n* = 8 mice, TLR4i: *n* = 9 mice. *P* values were calculated by one-tailed Mann-Whitney test. **i**, Expression of the indicated genes in ECs isolated from lungs harboring MDA231-LM2 metastases as in panel h; *n* = 3 or 4 mice. Mice harboring comparable lung metastatic loads, measured by *in vivo* bioluminescence imaging, were selected to analyze. *P* values were determined by one-way ANOVA with Dunnet’s multiple comparison test. * *P* < 0.05, ** *P* < 0.01, *** *P* < 0.001. For panels d, g and i, expression was determined by qPCR. **j,** Experimental outline of spontaneous lung metastasis in an orthotopic model where mice injected with MDA231-LM2 cells to fourth mammary fat pad were treated with TLR4i from day 14 to 24 after implantation of cancer cells. **k**, Spontaneous metastases in lungs of mice on day 24 in j. Left, representative examples of metastases (arrows) marked by expression of human vimentin by cancer cells. Right, quantification of lung metastasis area and nodule numbers based on human vimentin marker expression, vehicle: *n* = 15 mice, TLR4i: *n* = 12 mice. *P* values were calculated by one-tailed Mann-Whitney test. Scale bar, 200 μm. **l**, Model outlining the interaction between breast cancer cells and macrophages via TNC-TLR4 axis leading to macrophage activation and subsequent induction of the perivascular niche to produce pro-metastatic factors such as INHBB, OPG, LAMA1 and SCGB3A1. INHBB induces stem cell properties in breast cancer cells and OPG protects cancer cells from TRAIL-induced apoptosis at the metastatic site.

### Combination treatment with TLR4 inhibitor and anti-VEGF therapy impedes lung metastasis

As discussed earlier, we observed a discordance between EC genes regulated by VEGF or inflammatory cytokines from macrophages. This indicates that VEGF signaling and macrophages may induce two distinct responses in ECs during metastasis. We compared the gene expression profile of ECs isolated from metastatic lungs where the mice had been depleted of macrophages to the EC profile from mice treated with anti-VEGF antibody. Based on GSEA, the results indicate that macrophage-depended EC gene induction is particularly linked to inflammatory responses which associate with expression of GSP58, whereas VEGF-dependent induction results in stimulation of proliferation (Figure 8a). With these results in mind, we hypothesized that combining anti-TLR4 and anti-VEGF therapy may provide increased efficacy in suppressing metastasis. We treated xenograft and syngeneic metastasis-bearing mice with anti-TLR4 and anti-VEGF, individually or together (Figure 8b). The results showed that combined inhibition of TLR4 and VEGF provides improved efficacy in repression of metastasis compared to single treatments (Figure 8c,d). Together, our results describe the distinct endothelial activation properties of macrophage-mediated inflammation that induces production of vascular niche proteins and VEGF signaling that promotes proliferation of endothelial cells (Figure 8e). The results provide rationale to explore the combination of TLR4 inhibition with anti-VEGF therapy to effectively suppress the vascular functions in metastases.

**Figure 8.**
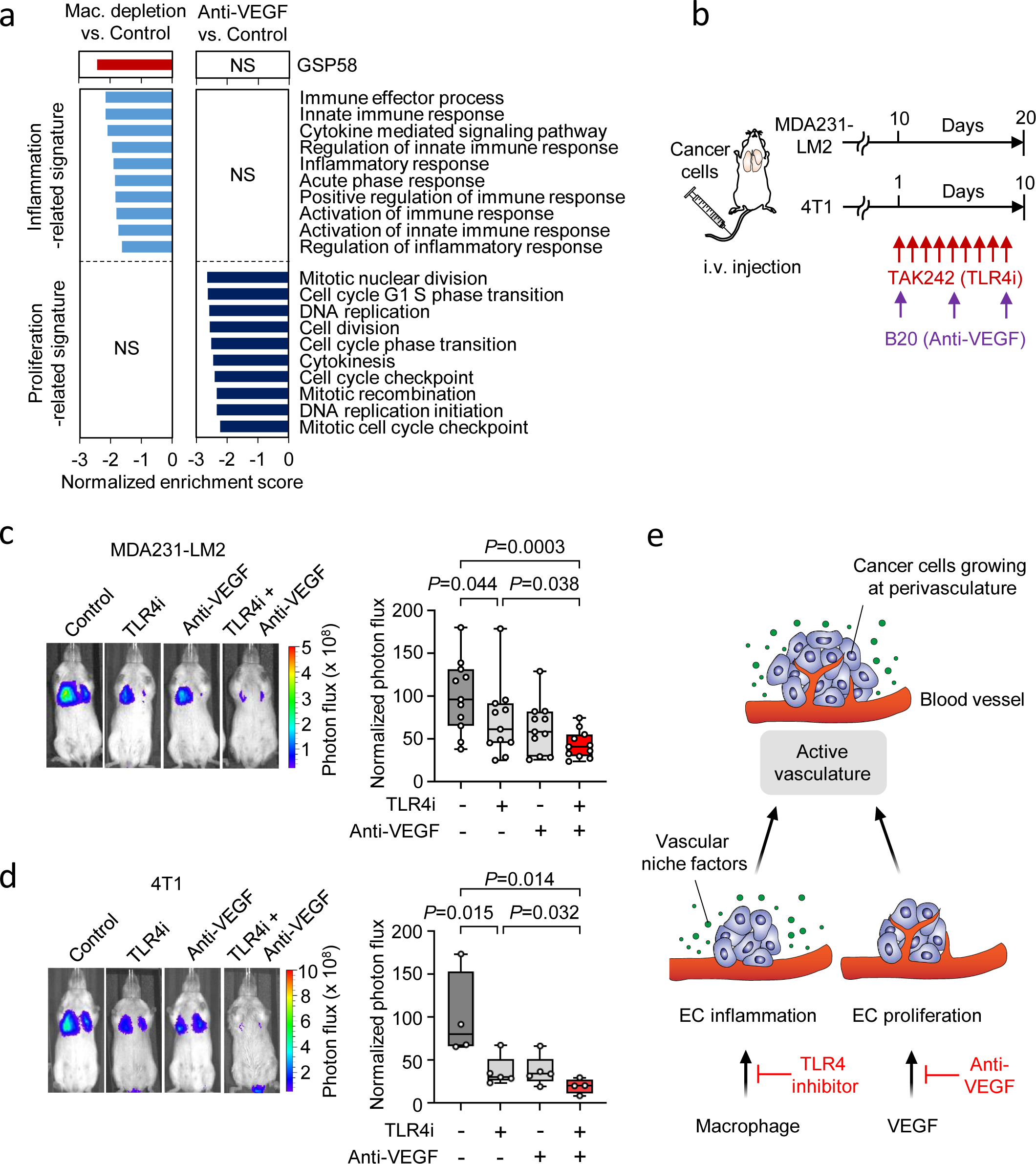
TLR4 inhibition and anti-VEGF therapy target different functions of the vascular niche with combination treatment impeding lung metastasis. **a**, GSEA of GSP58, inflammation- or proliferation-related signatures expressed in ECs from mice with lung metastases and depleted of macrophages (clodronate-liposomes) or treated with anti-VEGF therapy (B20.4.1.1). NS, not significant. **b**, Experimental outline where mice that have been intravenously injected with MDA231-LM2 or 4T1 cancer cells were treated with TLR4i or anti-VEGF as single treatments or together as a combination treatment. **c,d**, Bioluminescence analysis of metastatic colonization of the lung of mice injected with indicated cancer cells and treated as described in panel b. For both panels, Left shows representative bioluminescence images and right shows quantification of the metastatic colonization in the lungs based on the luminescence signal. MDA231-LM2, *n* = 11 mice; 4T1, *n* = 5 mice (single TLR4i and anti-VEGF treatment) and *n* = 4 mice (control and double treatment). *P* values were calculated by one-tailed Mann-Whitney test. **e**, Model depicting two regulatory arms of the vascular niche, where VEGF promotes proliferation of ECs and macrophages, activated by TNC-TLR4 signaling, promote inflammatory reaction in ECs and secretion of pro-metastatic factors of the vascular niche that are utilized by cancer cells.

## DISCUSSION

Cancer cells that disseminate to secondary organs must confront a microenvironment that is largely different from that of the primary site and thus likely to be unfavorable^29, 30^. Metastasis-initiating cancer cells and their progeny may be able to induce changes in the microenvironment at the secondary sites generating a niche that supports initiation and growth of metastasis^2, 3^. In this study, we revealed characteristic changes in metastasis-associating ECs in lungs, induced during metastatic colonization. Moreover, we demonstrated a crucial role of specific molecular crosstalk at perivascular sites between disseminated breast cancer cells, macrophages and vascular endothelium during lung metastasis.

Previous studies have shown that neo-angiogenesis is essential to fuel the growth of primary tumors^31^. This generated incentive to develop anti-angiogenic therapies that early on were primarily focused on targeting VEGF-mediating signaling^5, 31^. Whereas VEGF targeting as monotherapy has a promising effect in animal models and can promote overall survival of tumor bearing mice, this does not translate into overall survival in human cancer patients^4^. Anti-angiogenic therapy against VEGF combined with chemotherapy was approved against advanced metastatic breast cancer in 2007, but revoked four years later due to lack of improvement in overall survival^7^. Anti-angiogenic therapy has been shown to be effective in reducing formation of new vessels and may lead to cell death of immature tumor vessels and possible normalization^4^. However, established vessels are not eradicated. These mature vessels may still have an effect on metastatic colonization, by providing adhesion^9, 10^ and, as we showed, facilitating specific molecular crosstalk with disseminated cancer cells.

Growing evidence indicates that endothelial cells play a crucial role in intercellular communication during development, tissue homeostasis and disease. Transmembrane or secreted endothelial factors involved in cellular crosstalk have been termed angiocrine factors^32^. Previous studies have shown that angiocrine factor can play a role in cancer and influence disease progression. Endothelial cells confer cancer stem cell properties in breast cancer cells via Notch signaling, thus promoting aggressive behavior and mammary tumor growth^33^. Moreover, in mouse models for breast cancer metastasis to bone, angiocrine factors promote epithelial to mesenchymal transition and stem cell properties in cancer cells to further metastatic colonization^34^. Endothelial cells can also produce specific ECM proteins that regulate dormancy and possible transition to activation in disseminated breast cancer cells as well as potential resistance to therapeutic intervention^11, 35^. Our global transcriptomic analysis of metastasis-associated endothelial cells revealed specific gene expression patterns in metastasis-associated lung ECs that are linked to cell proliferation and migration and regulated by VEGF. However, our results show that inflammatory reaction of ECs, leading to production of numerous secreted proteins, GSP58, is largely independent of VEGF and plays an important role in supporting cancer cells in the metastatic lung. The findings suggest that anti-VEGF therapy may not be able to block angiocrine function of ECs during lung metastasis.

We identified a pro-metastatic crosstalk between disseminated breast cancer cells, macrophages and ECs in lung. Macrophages are activated by cancer cell-derived TNC and subsequently stimulate lung ECs to evoke the formation of a vascular niche. Notably, macrophage depletion suppressed not only the four vascular niche factors that we identified, but also substantial number of GSP58 that are induced in lung ECs. Our results provide insights into vascular regulation during metastasis, providing a distinction between VEGF-induced proliferation and macrophage-induced inflammatory responses in the regulation of vascular reactions. Association of macrophages and cancer cells at perivascular sites has been reported in mammary tumors in mouse models. Previous work, using multiphoton-based intravital imaging of mouse tumors, has revealed that increased vascular permeability and cancer cell-intravasation at vascular regions where cancer cells are joined by macrophages^36^. Our work shows that association of disseminated cancer cells, macrophages and ECs at the metastatic site is a crucial regulatory system of vascular niche formation where activated macrophages act as a trigger of angiocrine switch to promote the production of vascular niche factors in ECs.

We demonstrate that INHBB, LAMA1, SCGB3A1 and OPG are functional mediators of metastatic colonization and components of the vascular niche. We further analyzed the function of two of the factors, INHBB and OPG. INHBB is a subunit of proteins of the TGFbeta family such as activins and inhibins. In particular, activin B is formed by an INHBB homodimer. Activins are cytokines that are highly induced during wound healing, including the regeneration from vessel injury^37, 38^. Activin B has been shown to promote wound healing in mice via activation of RhoA-JNK signaling^39^. This is intriguing in light of our previous work showing that JNK signaling promotes stem cell properties in breast cancer cells^40^. Furthermore, our results are in line with the important role of activin in maintaining pluripotency in stem cells^41^. The second component of the vascular niche that we characterized was OPG that is recognized to have a pleiotropic function^19^. The best characterized function of OPG is its role as a decoy receptor of RANKL or TRAIL, antagonizing their function^42^. In addition to this role, OPG has been shown to promote survival in cells via integrin αVβ3-mediated NF-κB signaling^43^. Thus, the functions of OPG at the metastatic site may possibly be even broader than TRAIL neutralization.

The ECM protein TNC is recognized as a marked promoter of cancer progression in numerous malignancies^44, 45^. We revealed how cancer cells initiate activation of the vascular niche via TNC, produced by cancer cells, leading to macrophage activation and subsequent endothelial niche formation. Our results show that TNC, produced by cancer cells, activates perivascular macrophages through TLR4. We demonstrate that inhibition of TNC-TLR4 axis efficiently suppresses perivascular niche formation of breast cancer metastasis in the lungs. Notably, TNC-induced TLR4 activation has been shown to occur in the context of arthritis^28^. TNC is a glycoproteins within the ECM and is composed of multiple functional domains. The C-terminal domain called fibrinogen globe was previously shown to be the protein module that directly binds to and activates TLR4^28, 46^. In metastatic breast cancer cells, TNC expression has been shown to be curbed by the microRNA miR335 and induced by JNK signaling^40, 47^. In addition to exerting changes in the stroma, TNC expressed by breast cancer cells has been demonstrated to function in an autocrine manner, modulating Notch and Wnt signaling components to promote metastatic fitness of disseminated cancer cells Together, these results underscore the pleiotropic nature of TNC function in metastasis.

In conclusion, our findings delineate a complex crosstalk within growing metastasis in the lungs. The results show a crucial role for the vascular niche during metastatic progression and emphasize the role of the extracellular matrix in its regulation. These interactions in metastatic nodules may serve as viable targets when developing future therapies against metastatic disease.

## METHODS

### Cell culture

The breast cancer cell lines MDA-MB-231 (ATCC), MDA-MB-231-LM2 (provided by Joan Massagué, RRID:CVCL_5998)^12^ and 4T1 (ATCC) as well as the myeloid cell lines RAW264.7 (ATCC) and HL60 (ATCC) were cultured in Dulbecco’s Modified Eagle Medium (DMEM) GlutaMAX (ThermoFisher Scientific) supplemented with 10% fetal bovine serum (FBS, Gibco). SUM159 breast cancer cells (Asterand Bioscience) and the lung metastatic derivative SUM159-LM1 ^40^ were maintained in DMEM/F12 medium (ThermoFisher Scientific) with 5% FBS, 5 μg/ml human insulin (Sigma-Aldrich). E0771 cancer cells (CH3 BioSystems) were cultured in RPMI1640 medium with 10% FBS and 10 mM HEPES (Sigma-Aldrich). Primary human pulmonary endothelial cells (ECs) were purchased from Lonza and cultured in EBM-2 medium (Lonza) supplemented with EBM-2 bullet kit (Lonza). Human pulmonary EC line ST1.6R^48^ was provided by Ronald E. Unger and C. James Kirkpatrick and cultured on collagen type I (Corning) -coated dishes with M199 (ThermoFisher Scientific) supplemented with 20% FBS, 25 μg/ml endothelial cell growth supplement (Corning) and 25 μg/ml heparin (Sigma-Aldrich). For isolation and culture of bone marrow-derived macrophages, bone marrow cells were obtained from 8-10 weeks old BALB/c mice and cultured for 7 days in DMEM supplemented with 10% FBS and 20 μg/ml macrophage-colony stimulating factor (M-CSF, Peprotech). All medium contained 50 U/ml penicillin and 50 μg/ml streptomycin (Sigma-Aldrich).

### Mouse experiments

Female non-obese diabetic-severe combined immunodeficiency gamma^null^ (NSG, The Jackson Laboratory), BALB/c (Janvier Labs or Envigo) or C57BL/6 (wild-type) 6-12 week old mice were used for in vivo studies. Mice were housed in individually ventilated cages with temperature and humidity control and under 12-12 h light-dark cycle. All experiments with mice were conducted according to the German legal regulations, and protocols were approved by the governmental review board of the state of Baden-Wuerttemberg, Regierungspraesidium Karlsruhe.

For lung colonization assay, 25,000-500,000 cells of cancer cells, suspended in 100 μl of phosphate-buffered saline (PBS) were injected via the tail vein. Human breast cancer cell line MDA231, MDA231-LM2, SUM159 and SUM159-LM1, and mouse mammary tumor cell line 4T1 and E0771 were transduced with a triple reporter expressing herpes simplex virus thymidine kinase 1, green fluorescent proteins (GFP), and firefly luciferase genes^49^, enabling bioluminescence imaging. To monitor lung metastatic progression, mice were intraperitoneally injected with 150 mg/kg of D-Luciferin (Biosynth), anesthetized using isoflurane (Orion) and imaged with IVIS Spectrum Xenogen machine (Caliper Life Sciences). Bioluminescence images were analyzed using Living Image software, version 4.4 (Caliper Life Sciences).

For spontaneous lung metastasis assay in an orthotopic mouse model, 500,000 MDA231-LM2 breast cancer cells were suspended in a 1:1 (vol/vol) mixture of Growth Factors-Reduced Matrigel (Corning) and PBS, and injected bilaterally to the fourth mammary fat pads, in a volume of 50 μl. Primary tumor volume was measured using digital caliper and calculated with the following formula: volume = length x width^2^ x 0.52. Mice were sacrificed and metastatic burden in the lungs was analyzed by immunohistochemistry 24 days post cancer cell implantation.

### Drug treatment *in vivo*

For the inhibition of vascular endothelial cell growth factor (VEGF) signaling *in vivo*, mice harboring lung metastases were intraperitoneally injected with anti-human/mouse VEGF-A neutralizing antibody B20.4.1.1 (5 or 10 mg/kg, twice in a week, provided by Genentech)^14^ or anti-mouse VEGFR2 blocking antibody DC101 (40 mg/kg, on day 21 and 24 after the injection of cancer cells, BioXcell). For macrophage-depletion, mice were intravenously injected with clodronate-liposome (100-150 μl per mouse, every other day, Liposoma BV). Short-term experiments were performed by treatment with clodronate-liposome on day 14, 16 and 18 after injection of MDA231-LM2 cells, or on day 5, 7 and 9 after injection of 4T1 cells. Whereas for long-term experiments, mice were treated with clodronate-liposome for the duration of the experiment, starting 1 day before tail vein injection of cancer cells. PBS-loaded liposome (Liposoma BV) was used as a control treatment. For neutrophil-depletion, anti-Ly6G antibody (1A8, 400 μg per mouse, BioXcell) was intraperitoneally injected on day 5, 7 and 9 after the tail vein injection of 4T1 cells. For toll-like receptor 4 (TLR4) inhibition, TAK-242 (10 mg/kg, once a day, Merck Millipore) was intraperitoneally injected to mice harboring lung metastases from day 10 to 20 (MDA231-LM2 metastases) or from day 1 to 10 (4T1 metastases), following intravenous injection of cancer cells. To inhibit TLR4 in a spontaneous metastasis model, mice were treated with TAK-242 from day 14 to 24 after orthotopic injection of MDA231-LM2 breast cancer cells. Combination of TLR4 inhibition and VEGF inhibition was performed by intraperitoneal injection of TAK-242 (10 mg/kg, once a day) and B20.4.1.1 (10 mg/kg, twice a week) from day 10 to 20 for lung colonization of MDA231-LM2 cells, and from day 1 to 10 for 4T1 cancer cells.

### Fluorescence-activated cell sorting

Mouse lung ECs were isolated by fluorescence-activated cell sorting (FACS). Lungs with or without metastasis were digested using 0.5% Collagenase type III (Pan Biotech), 1% Dispase II (Gibco) and 30 μg/ml DNase I in PBS for 45 min at 37°C. Single cell suspensions were generated by repeated pipetting in FACS buffer (PBS containing 2% FBS and 2 mM EDTA), and filtering through 70 μm nylon filters. Following centrifugation, the cell pellets were suspended in ACK Lysing Buffer (Lonza) and incubated for 5 min at room temperature (RT) to remove red blood cells. After washing with FACS buffer, cells were suspended with FcR blocking reagent (Miltenyi Bio Tec) diluted in FACS buffer (1:10) and incubated for 10 min on ice. Cells were then stained for 30 min on ice with the following antibodies: phycoerythrin (PE)-conjugated anti-CD45 (30-F11, 1:3000, eBioscience), PE-conjugated anti-CD11b (M1/70, 1:3000, BD Biosciences), PE-conjugated anti-CD326 (G8.8, 1:250, eBioscience), Allophycocyanin (APC)-conjugated anti-CD140a (APA5, 1:50, eBioscience), APC-conjugated anti-CD140b (APB5, 1:50, eBioscience) and PE-Cyanine7 (Cy7)-conjugated anti-CD31 (390, 1:500, eBioscience). After washing with FACS buffer, cell sorting was performed with BD FACSAria I or FACSAria II machines (Becton Dickinson). GFP (cancer cells)-negative, PE-negative, APC-negative and PE-Cy7-positive population was isolated as lung EC fraction. Fibroblasts (CD140a and CD140b-positive), bone marrow-derived cells (CD45-positive) and epithelial cells (CD326-positive) were also isolated and the expression of vascular niche factor candidates in these cell types was analyzed by quantitative polymerase chain reaction (qPCR). To analyze populations of macrophages and neutrophils in lung, mouse lungs were digested as described above, and cells incubated for 30 min on ice with following antibodies: PE-conjugated anti-CD11b (M1/70, 1:3000, BD Biosciences), APC-conjugated anti-F4/80 (for macrophages, BM8, 1:400, eBioscience) or APC-conjugated anti-Ly6G (for neutrophils, RB6-8C5, 1:2000, Invitrogen). FACS data obtained by these experiments were analyzed using FlowJo V10 software.

### Analysis of gene expression profiles

Gene expression profiles of ECs sorted from mouse lung were generated by transcriptomic analysis using Affymetrix GeneChip Mouse Genome 430 2.0 or Human Genome U133 Plus 2.0 Arrays according to the manufacturer’s protocol. Raw CEL-files were RMA-normalized and clustered with principle components using Chipster (version 3.8.0) or R (version 3.5). The differential gene expression analysis was performed by two-group comparisons using empirical Bayes test with Benjamini-Hochberg correction of *P* values. Gene Ontology (GO) analysis was conducted with the Database for Annotation, Visualization, and Integrated Discovery (DAVID)^50^, and EC secretome gene set was generated utilizing GO: 0005576 (GO term: extracellular region). Heatmap images were produced by MeV 4.9.0 software and volcano plot was generated with R version 3.5. Gene set enrichment analysis (GSEA) was performed with Molecular Signatures Database (MSigDB) from the Broad Institute^51^. Nominal *P* values were calculated based on random gene set permutations with Benjamini-Hochberg correction and FDR < 0.25 were regarded as statistically significant. For the analysis of gene expression signatures with violin plots, genes of specific gene sets described in each figure legends were z-scored, and the average z-score from 3 biological replicates was calculated and plotted in figures. Gene expression profiles of distant metastasis from breast cancer patients (GSE14020 or the lung metastasis patients of GSE14018) were RMA-normalized using R version 3.5. The hierarchal clustering was performed based on the expression of EC-secretome genes with heatmap function in the R statistical package, and obtained clusters were used in GSEA and survival analysis. Pearson’s correlation analysis of vascular niche genes with *Cdh5*, *Tnc*, and signature of classically activated macrophages (Orecchioni, M. et al., 2019) was conducted with the GSE14020 dataset using GraphPad Prism 8.

Survival analysis of breast cancer patients with GSP58-high and -low clusters was performed using datasets of lung metastasis of GSE14018. Datasets from the Kaplan-Meier plotter (KM plotter, https://kmplot.com/analysis/) were utilized to analyze the association of *Inhbb*, *Lama1, Sgb3a1* and *Opg* with survival of ER-negative breast cancer patients. Datasets used in KM plotter were following: E-MTAB-365, GSE16716, GSE17907, GSE19615, GSE20271, GSE2034, GSE20711, GSE21653, GSE2603, GSE26971, GSE2990, GSE31519, GSE3494, GSE37946, GSE42568, GSE45255, GSE4611, GSE5327 and GSE7390 for relapse-free survival; GSE16716, GSE20271, GSE20711, GSE3494, GSE37946, GSE42568, GSE45255 and GSE7390 for overall survival.

### Immunohistological analysis

Dissected mouse lungs were fixed with 10% buffered formalin for 4-12 h at 4°C, incubated with 30% sucrose/PBS overnight, and embedded in O.C.T. compound (Sakura Finetek). Cryostat sections with 8 μm thickness were prepared with Microm HM-525 cryotome (Thermo Fisher Scientific).

For immunofluorescence staining, sections were washed with PBS, blocked with blocking solution consisting of 0.5% Blocking Reagent (Perkinelmer), 0.1 M Tris-HCl (pH.7.5) and 0.15 M NaCl for 1h at RT, and incubated with anti-GFP (ab290 or ab13970, 1:1000 dilution, abcam), anti-CD31 (Mec13.3, 1:100 dilution, BD Pharmingen, or ab28364, 1:50 dilution, abcam), anti-cleaved caspase 3 (D3E9, 1:250 dilution, Cell Signaling), anti-F4/80 (BM8, 1:100 dilution, Invitrogen), anti-TNC (BC-24, 1:4000 dilution, ThermoFisher Scientific) or anti-TNFα (AF-410, R&D Systems, 10 μg/ml) at 4°C overnight. After being washed with 0.05% Tween 20/PBS, sections were stained with fluorescence-conjugated secondary antibodies and DAPI (BioLegend) for 1h at RT. Sections were then washed with 0.05% Tween 20/PBS and mounted with Fluoromount-G (SouthernBiotech). Images were obtained with Cell observer microscope (Zeiss) and analyzed with FIJI (ImageJ) or ZEN imaging software (Zeiss).

For vimentin staining, sections were rehydrated with decreasing concentrations of ethanol, quenched with 3 % hydrogen peroxide, and antigen retrieval was carried out at 100°C for 20 min with citrate buffer (pH6.0, Vector Laboratories). Sections were blocked with 0.1% bovine serum albumin (BSA) containing 0.1% Triton-X100 for 2 h at RT, followed by incubation with anti-vimentin antibody (SRL33, 1:400 dilution, Leica Biosystems). Biotinylated anti-mouse IgG secondary antibody and ABC avidin-biotin-DAB detection kit (Vector laboratories) were used for signal detection according to manufacturer’s instructions. Sections were counterstained with Mayer’s hematoxylin solution (Sigma-Aldrich) for 1 min, dehydrated using increasing concentrations of ethanol and mounted using Cytoseal XYL (ThermoFisher Scientific). Images were obtained with Cell observer microscope (Zeiss) and analyzed with FIJI (ImageJ) or ZEN imaging software (Zeiss).

### Oncosphere formation

SUM159-LM1 cells or MDA231-LM2 cells were suspended in HuMEC/0.1% BSA and treated with EC-derived conditioned media (CM) (diluted 2-5 times in the final medium) or 50 ng/ml of recombinant activin B (R&D systems). 2,500 cells/well of SUM159-LM1 cells or 5,000 cells/well of MDA231-LM2 cells were seeded on 96-well ultra-low attachment plate (Corning), and cultured for 7 days. To prepare human lung EC-derived CM, confluent ST1.6R cells were cultured in HuMEC medium (Invitrogen) supplemented with 0.1% BSA for 24h, and the culture media was collected and filtered with 0.45 μm pore size of syringe-filter (Techno Plastic Products). The number of oncospheres per well was determined by using a Zeiss Primovert microscope (Zeiss).

### Lentivirus-mediated gene overexpression and knockdown

For overexpression of vascular niche genes, full length cDNAs of human INHBB and OPG were generated by PCR with a template total cDNA obtained from human pulmonary EC line ST1.6R. Human SCGB3A1 cDNA was synthesized as GeneArt Strings DNA Fragments (Invitrogen). Obtained cDNAs were sub-cloned into pLVX-Puro lentiviral expression vector (Clontech) and transfected to HEK293T cells together with packaging plasmids psPAX2 and pMD2G using Lipofectamine 2000 (Invitrogen). Viral supernatants were collected after 48 h, and used to infect cancer cells in the presence of 8 μg/ml polybrene (Sigma-Aldrich). Infected cells were selected with 2 μg/ml puromycin (Invitrogen) for 7 days, and gene overexpression was confirmed by qPCR. Human LAMA1 cDNA was purchased from Promega and sub-cloned into pLVX-Tet-On Advanced vector (Clontech). Using the combination with pLVX-Tight-puro vector (Clontech), doxycycline-mediated inducible expression clone of cancer cells were generated by screening with 2 μg/ml puromycin and 600 μg/ml Zeocin (ThermoFisher Scientific).

For knockdown experiments, The shERWOOD algorithm^52^ was utilized to design shRNAs, and following sequences of oligonucleotides were used: 5’-TGCTGTTGACAGTGAGCGAAACCATCAAACTTCATGATCATAGTGAAGCCACAGATGTATGATCATGAAGTTTGATGGTTGTGCCTACTGCCTCGGA-3’ for human DR4 shRNA#1, 5’-TGCTGTTGACAGTGAGCGACTCCCATGTACAGCTTGTAAATAGTGAAGCCACAGATGTATTTACAAGCTGTACATGGGAGGTGCCTACTGCCTCGGA-3’ for human DR4 shRNA#2, 5’-TGCTGTTGACAGTGAGCGCCTCACTGGAATGACCTCCTTATAGTGAAGCCACAGATGTATAAGGAGGTCATTCCAGTGAGTTGCCTACTGCCTCGGA-3’ for human DR5 shRNA#1, 5’-TGCTGTTGACAGTGAGCGACAAGACCCTTGTGCTCGTTGATAGTGAAGCCACAGATGTATCAACGAGCACAAGGGTCTTGGTGCCTACTGCCTCGGA-3’ for human DR5 shRNA#2, and 5’-TGCTGTTGACAGTGAGCGACCAGCCTGTGGCTTTACCTGATAGTGAAGCCACAGATGTATCAGGTAAAGCCACAGGCTGGCTGCCTACTGCCTCGGA-3’ for human INHBB shRNA. Human TNC was targeted as previously described^26^. Oligonucleotides were cloned into lentivirus vector StdTomatoEP or StdTomatoEZ, and transfected to HEK293T cells with psPAX2 and pMD2G using Lipofectamin 2000. Generated lentiviruses were used for the infection of MDA231-LM2 or ST1.6R and knock down efficiency was assessed by qPCR after the selection of cells with 2 μg/ml puromycin and/or 600 μg/ml Zeocin for 5-7 days.

### Real time-qPCR

Total RNA was extracted using RNeasy Mini kit (Qiagen) or Arcturus PicoPure RNA isolation kit (Applied Biosystems), and reverse transcription was performed with High-Capacity cDNA Reverse Transcription kit (Applied Biosystems) according to the manufacture’s instructions. Real time-qPCR was performed with SYBR Green gene expression assay (Applied Biosystems) using ViiA 7 Real-Time PCR System (Applied Biosystems). Primer pairs listed in Supplementary table 3 were used.

### Stimulation of ECs with conditioned media

For preparation of cancer cell-derived CM, MDA231-LM2 cells were cultured in DMEM/1% FBS for 24 h and medium was collected as a cancer cell-CM. For production of CM from HL60 cells, undifferentiated HL60 cells, cultured in RPMI1640/10% FBS, were stimulated with 1.25% dimethyl sulfoxide (DMSO, Sigma-Aldrich) and 1 μM all-trans retinoic acid (Sigma-Aldrich) to induce the differentiation towards neutrophil lineage. After 2 days of culture, cells were stimulated again with same stimulus, and further cultured for 2 days. Differentiated HL60 cells were then washed twice with PBS and cultured with RPMI1640/1% FBS for 24 h. Supernatants were collected and used for stimulation of ECs. For preparing the CM from a macrophage cell line, RAW264.7 cells cultured in DMEM/10% FBS were stimulated with 10 nM PMA or 24 h. Following PBS washing (twice), activated RAW264.7 cells were cultured in DMEM/1% FBS for 24 h and supernatant collected.

Primary HPMECs or ST1.6R cells were seeded on collagen I-coated dishes in EBM2 medium with 1% FBS or M199 containing 1% FBS, respectively, and cultured for 1 day. Cells were then stimulated with CMs prepared as above for 24 h and gene expression changes were analyzed by qPCR.

### Macrophage activation with TNC

Bone marrow-derived macrophages were cultured in DMEM/1% FBS containing 20 μg/ml M-CSF overnight. Cells were then treated with 3 μM TAK-242 (Merck Millipore) or vehicle control (0.1% DMSO in final) for 1 h, and 2 μg/ml of recombinant TNC (Merck Millipore) added to the medium. After 6 hours of stimulation, cells were collected and gene expression was assessed by qPCR.

### Stimulation of ECs with TNC

Primary HPMECs in EBM2 medium or ST1.6R cells in DMEM/1% FBS were cultured overnight and stimulated with 2 μg/ml of recombinant TNC (Merck Millipore) for 24 h. Cells were then collected and gene expression was analyzed by qPCR.

### Detection of TRAIL-induced apoptosis

MDA231-LM2 cells in DMEM with 10% FBS were cultured with or without incremental dosages (10-200 ng/ml) of recombinant OPG (R&D Systems) for 30 min, and treated with 50 ng/ml of recombinant TRAIL (R&D Systems) for 4 hrs. Cells were washed once with PBS, and lysed with RIPA buffer supplemented with 1 x HALT protease and phosphatase inhibitor cocktail (ThermoFisher Scientific).

Western blots were carried out as previously described^40^. Sources of primary antibodies used are as follows: anti-Cleaved caspase 3 (#9664, 1:500 dilution, Cell signaling), anti-Caspase 3 (#9662, 1:1000 dilution, Cell signaling), anti-Vinculin (#4650, 1:1000 dilution, Cell Signaling). Following the incubation with a primary antibody, membranes were washed and probed with horse radish peroxidase-conjugated IgG (1:10,000, Leica) and exposed to X-ray films (Fuji-film) with Clarity Western ECL Substrate (Bio-Rad).

### Statistical analysis

Statistical analyses were performed as described in figure legends. *P* < 0.05 was considered as significant and statistical tests for *in vitro* experiments were two-tailed unless otherwise indicated. All functional *in vivo* experiments were based on substantial *in vitro* results that indicated one-directional effect, and thus one-tailed tests were adopted for these experiments. Statistics for averaged Z-score of gene signatures were conducted with one-way ANOVA with Fisher’s least significant difference test unless otherwise indicated. For Kaplan-Meier analyses in breast cancer patients, statistical differences in survival curves were calculated by log rank (Mantel-Cox) test.

The GSE14020 gene expression dataset was used to study the correlation between vascular niche genes and *Cdh5*, *Tnc*, or signature of activated macrophages in a cohort of 65 metastasis samples from breast cancer patients. Gene expression values for each gene within individuals were associated in a correlation matrix. Then, Pearson correlation coefficient (r) and *P* value were calculated for each comparison. Statistical significance of gene expression data from microarray were calculated using the software Chipster. Statistical analyses of gene ontology and gene set enrichment were conducted using DAVID^50^, and GSEA^51^, respectively. For GSEA, an FDR < 0.25 was considered statistically significant. All other statistical analyses were performed using GraphPad Prism version 8 for Windows.

## ACKNOWLEDGEMENTS

We thank members of the Oskarsson lab for critical reading of the manuscript. We are grateful to the microarray unit of the DKFZ Genomics and Proteomics Core Facility, the Central Animal Laboratory, the Flow Cytometry Core Facility and the Light Microscopy unit of the Imaging and Cytometry Core Facility for advice and technical assistance. We thank Joan Massagué, Ronald E. Unger and C. James Kirkpatrick for sharing cell lines. Genentech kindly provided the B20.4.1.1 antibody against VEGF, via MTA program. T.H. was supported by the International Tenure Track program of University of Tsukuba and an overseas research fellowship from the Uehara memorial foundation. M.P. was supported by a scholarship from the Helmholtz International Graduate School for Cancer Research. This work was funded by the Dietmar Hopp Foundation.

## AUTHOR CONTRIBUTION

T.H. and T.O. designed experiments, analyzed data, and wrote the manuscript. T.H. performed experiments. M.P. and J.I.R. helped with analysis of gene expression profiles and immunohistological analyses. J.M. assisted with mouse experiments and K.D. supported qPCR analysis. A.D. and A.R. contributed to experimental design. T.O. supervised the research.

## COMPETING INTERESTS

The authors declare no competing financial interests

## SUPPLEMENTARY FIGURE LEGENDS

**Supplementary Figure 1.**
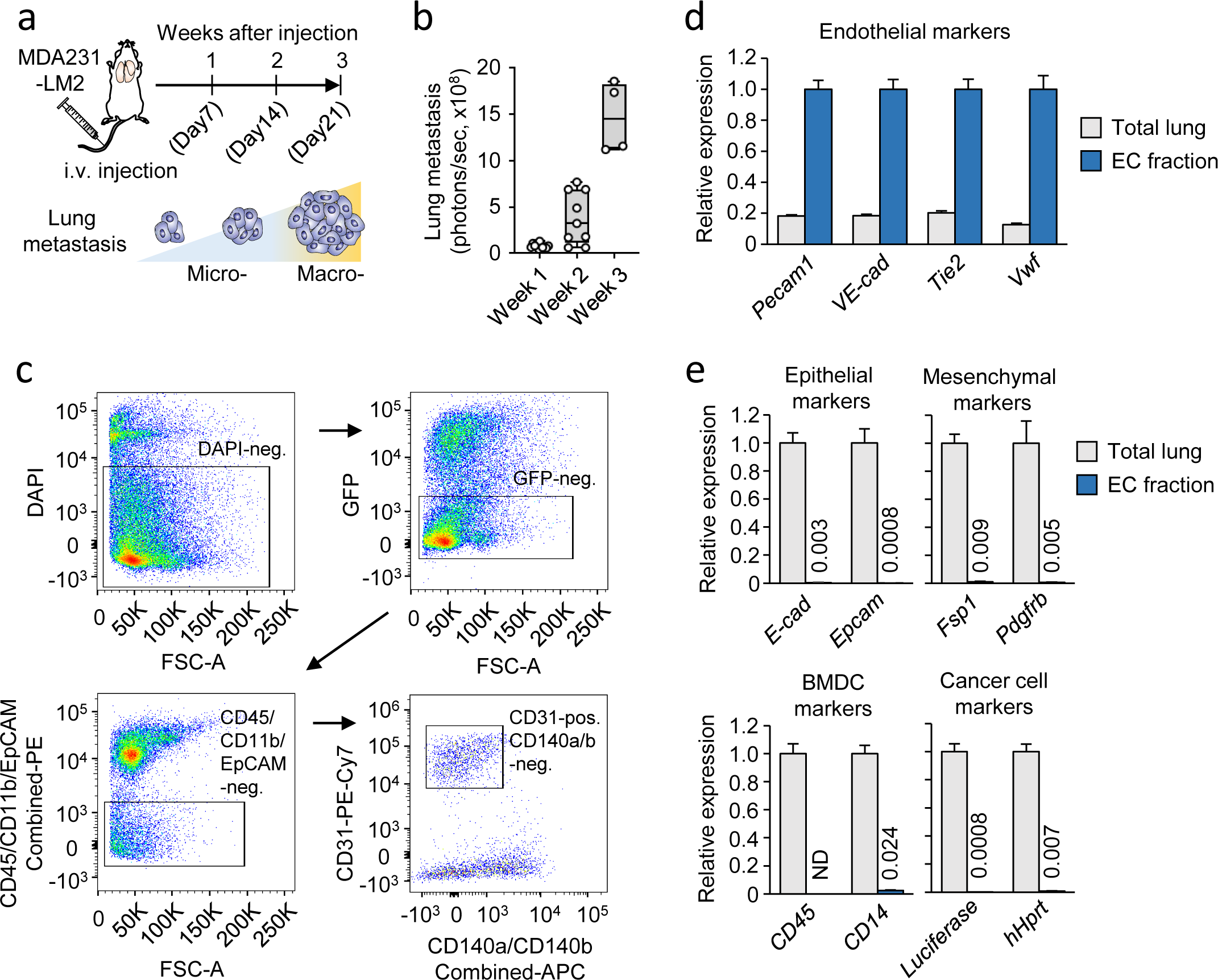
**a**, Schematic of the experimental outline in Figure 1. MDA231-LM2 cells were injected intravenously into NSG mice and lung metastasis was analyzed at week 1, week 2 and week 3 post injection. **b**, Lung colonization determined by bioluminescence in mice at indicated time points; *n* = 11 for week 1, *n* = 7 for week 2, *n* = 4 for week 3. **c**, Representative FACS plots showing isolation of ECs from lungs with metastasis. **d-e**, Expression of cell type-specific markers in total lung cells containing MDA231-LM2 metastases and sorted lung EC fractions. Endothelial cell makers (d), epithelial cell-, mesenchymal cell-, bone marrow-derived cell-(BMDC), and human cancer cell markers (e); numbers are expression levels relative to control; *n* = 3 experiments. ND, not detected.

**Supplementary Figure 2.**
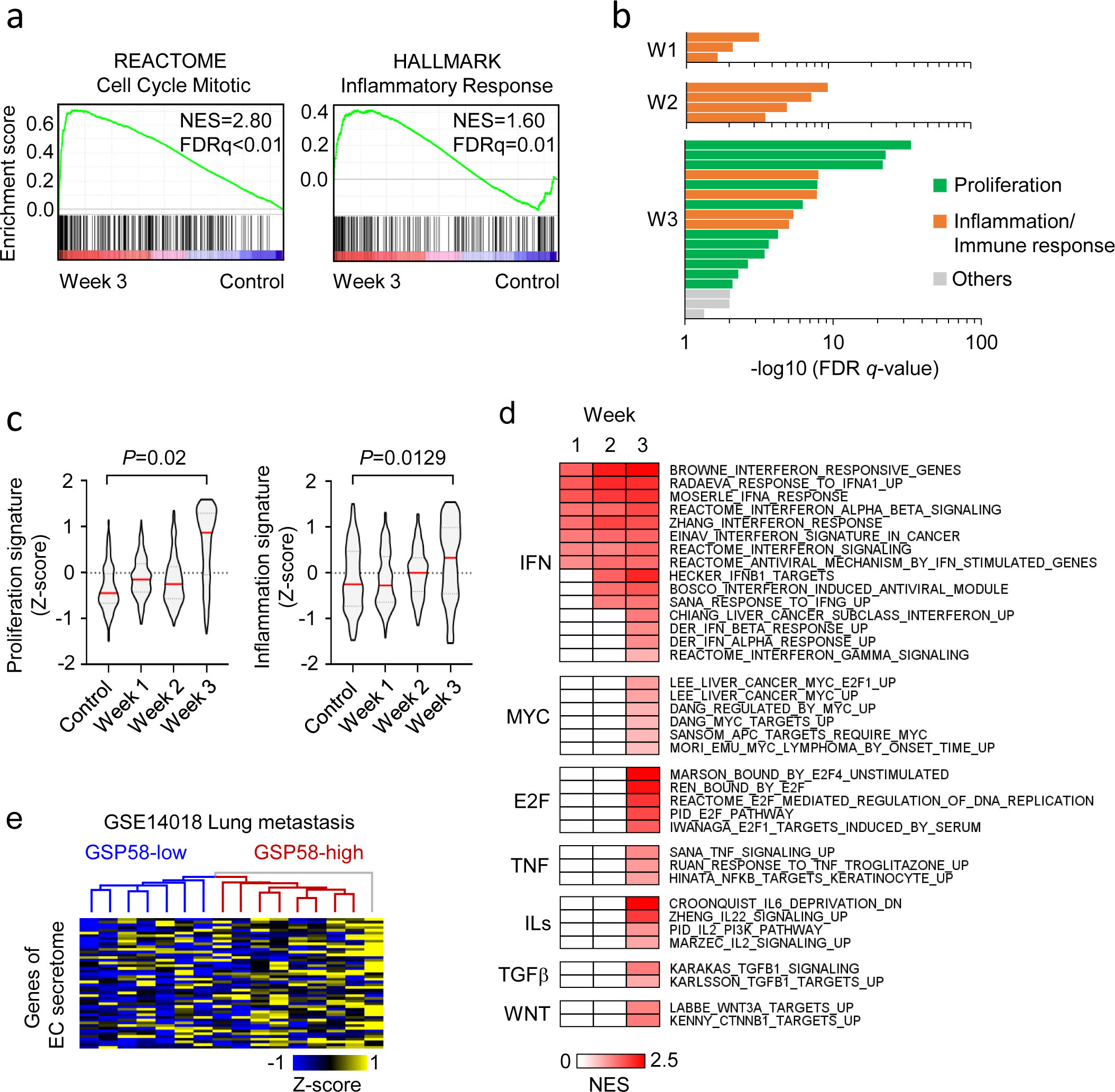
**a**, GSEA plots for proliferation signature (“Cell cycle mitotic” in REACTOME, left) and inflammation signature (“Inflammatory response” in Hallmark of MSigDB, right) in lung ECs isolated at week 3 compared to that of healthy control. FDR was determined from *P* values calculated by random permutation test. **b**, Gene ontology (GO) analysis of upregulated genes (FDR < 0.05, log2FC > 1) within metastatic EC transcriptome at indicated time points. Biological processes with FDR < 0.05 are shown. The specific GO terms in graph are listed in Supplementary table 1. **c**, Violin plot analysis of Z-scores from gene signatures for proliferation (Cell cycle mitotic, REACTOME, left) and inflammation (Inflammatory response, Hallmark in MSigDB, right). Z-scores were calculated from gene expression profiles of lung ECs isolated from healthy control lungs and metastatic lungs at week 1, 2 and 3. *P* values were determined by unpaired two-tailed t-test, using averaged Z-scores of 3 biological replicates. **d**, GSEA of signaling pathways (C2 collection of MSigDB) enriched in ECs at indicated time points. Signatures with nominal *P* < 0.05 and FDR < 0.25 were considered as significant. NES, normalized enrichment score. **e**, Hierarchical clustering of lung metastasis samples from breast cancer patients (GSE14018, 16 patients) according to the expression of GSP58.

**Supplementary Figure 3.**
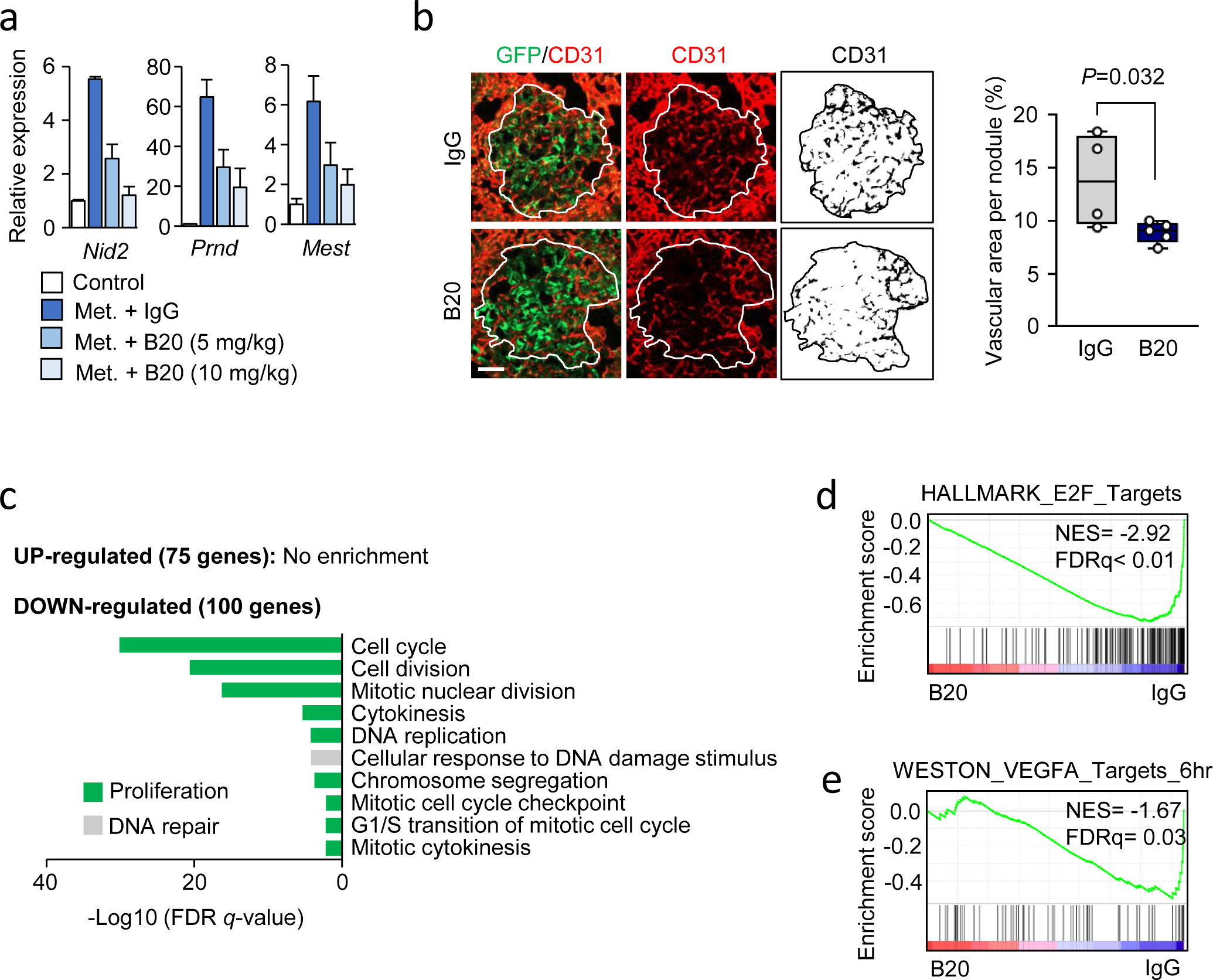
**a**, Relative expression of VEGF target genes in lung ECs isolated from healthy control mice or mice with lung metastasis (at week 3) treated with IgG or anti-VEGFA antibody (B20); *n* = 3 experiments. **b**, Immunofluorescence analysis of ECs (CD31, red) in metastatic nodules in lungs at week 3 from mice treated with control IgG or B20 antibodies. MDA231-LM2 cancer cells are green (GFP). Representative images (left) and quantification of vascular area per nodule (right) are shown. Scale bar, 50 μm. *P* value was determined by one-tailed Mann-Whitney test; n ≥ 4 mice. **c**, GO analysis of differentially regulated genes (*P* < 0.05, log2FC > ± 0.5) in lung ECs by the *in vivo* treatment with B20. GO terms of biological processes enriched with FDR < 0.05 are shown. **d-e**, GSEA plots of “E2F targets” (Hallmark in MSigDB) (d) and “VEGF targets” ^58^ (e) comparing lung ECs from mice treated with IgG or B20, as in Figure 2f. NES, normalized enrichment score.

**Supplementary Figure 4.**
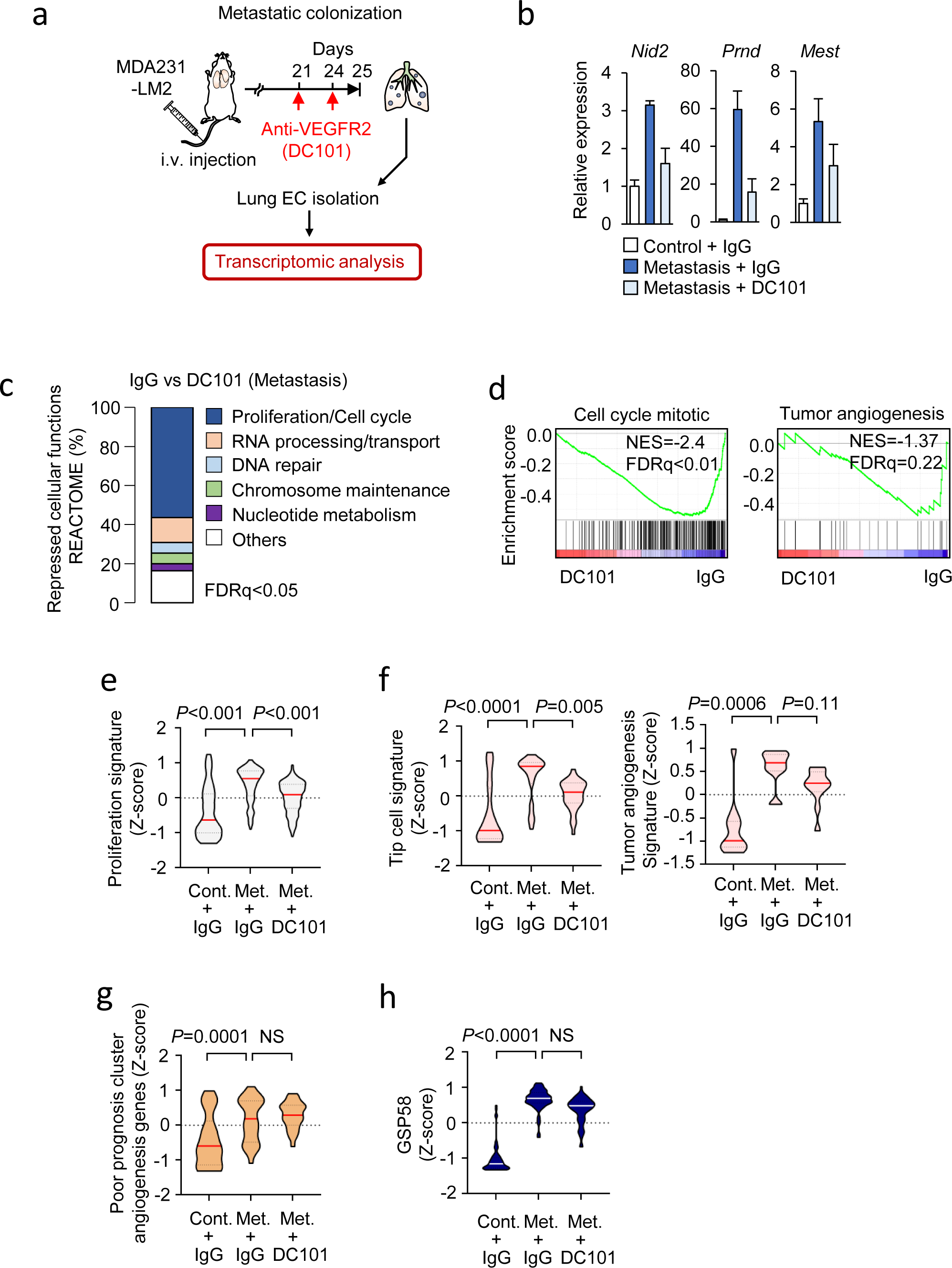
**a**, Schematic depicting experimental outline for metastatic colonization of the lung to analyze the effect of anti-VEGFR2 antibody treatment on metastasis-associated lung ECs. MDA231-LM2 cells were injected intravenously into mice, followed by treatment with anti-VEGFR2 antibody (DC101), or control IgG before dissection of the lung. ECs were isolated from lungs with metastases and transcriptomic analysis was performed. Mice harboring comparable metastatic loads in lungs, measured by *in vivo* bioluminescence imaging, were selected for analysis. **b**, Relative expression of VEGF target genes in lung ECs isolated from healthy control mice or mice with lung metastasis treated with IgG or DC101; *n* = 3 experiments. **c**, Overview of REACTOME pathway genes down-regulated by DC101. Pathways with FDR < 0.05 are shown. **d**, Repression of gene signatures of “Cell cycle mitotic” (REACTOME, left) and “Tumor angiogenesis” (MSigDB, right) in lung ECs with DC101 treatment compared to IgG control treatment. NES, normalized enrichment score. **e-h**, Z-scores of genes from indicated signatures expressed in ECs from lungs of mice under indicated conditions. Signatures: proliferation (“Cell cycle mitotic” in REACTOME) (e), tip cells^54^ (f, left) and tumor angiogenesis^53^ (f, right), poor prognosis angiogenesis genes^56^ (g) and GSP58 (h). *P* values were determined from Z-scores of genes within the signatures (e) or averaged Z-scores of 3 biological replicates (f-h) by one-way ANOVA with Dunnet’s multiple comparison test; *n* = 3 for each group. NS, not significant.

**Supplementary Figure 5.**
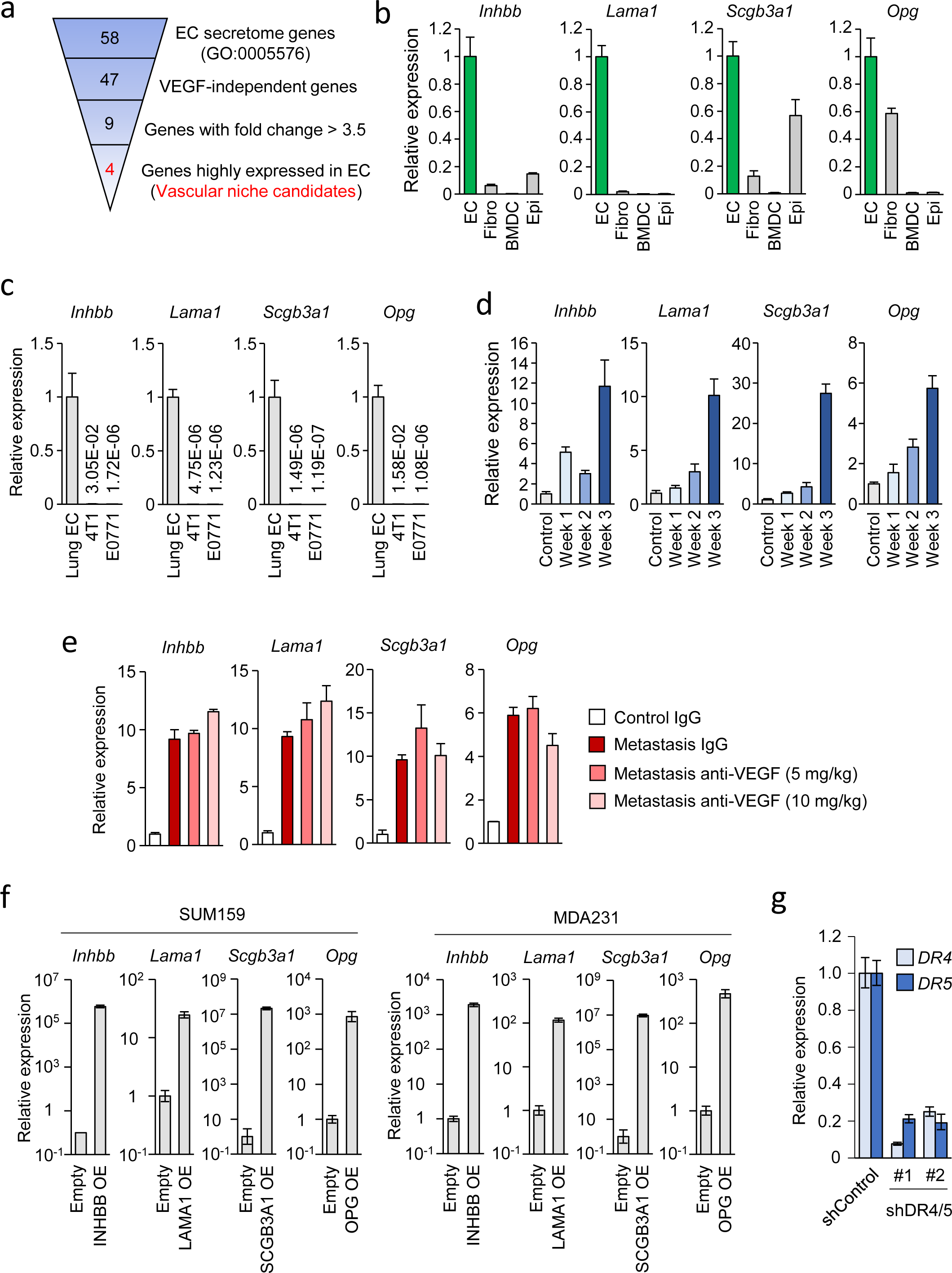
**a**, Overview of the process used to narrow down the gene candidates functioning within the pro-metastatic vascular niche during breast cancer metastasis to lung. **b**, Relative expression of the four vascular niche factors in lung ECs, fibroblasts (Fibro), bone marrow-derived cells (BMDC) and epithelial cells (Epi) isolated from lung with metastasis at week3, as in Figure 3a. **c**, Relative expression of the niche factors in lung ECs and the mouse mammary tumor cells 4T1 and E0771. **d**, Expression kinetics of niche components during lung metastatic progression. **e**, Relative expression of the four vascular niche factors in lung ECs isolated from lungs of healthy control mice or mice harboring metastasis treated with IgG or anti-VEGF antibody (B20). **f**, Expression of vascular niche factors in SUM159 (left) or MDA231 (right) cancer cells transduced with control vector or vectors overexpressing cDNA of each gene. **g**, Expression of TRAIL activated death receptors 4/5 (DR4/5) in MDA231 cells transduced with shControl or shDR4/5 (#1 and #2 are independent hairpins).

**Supplementary Figure 6.**
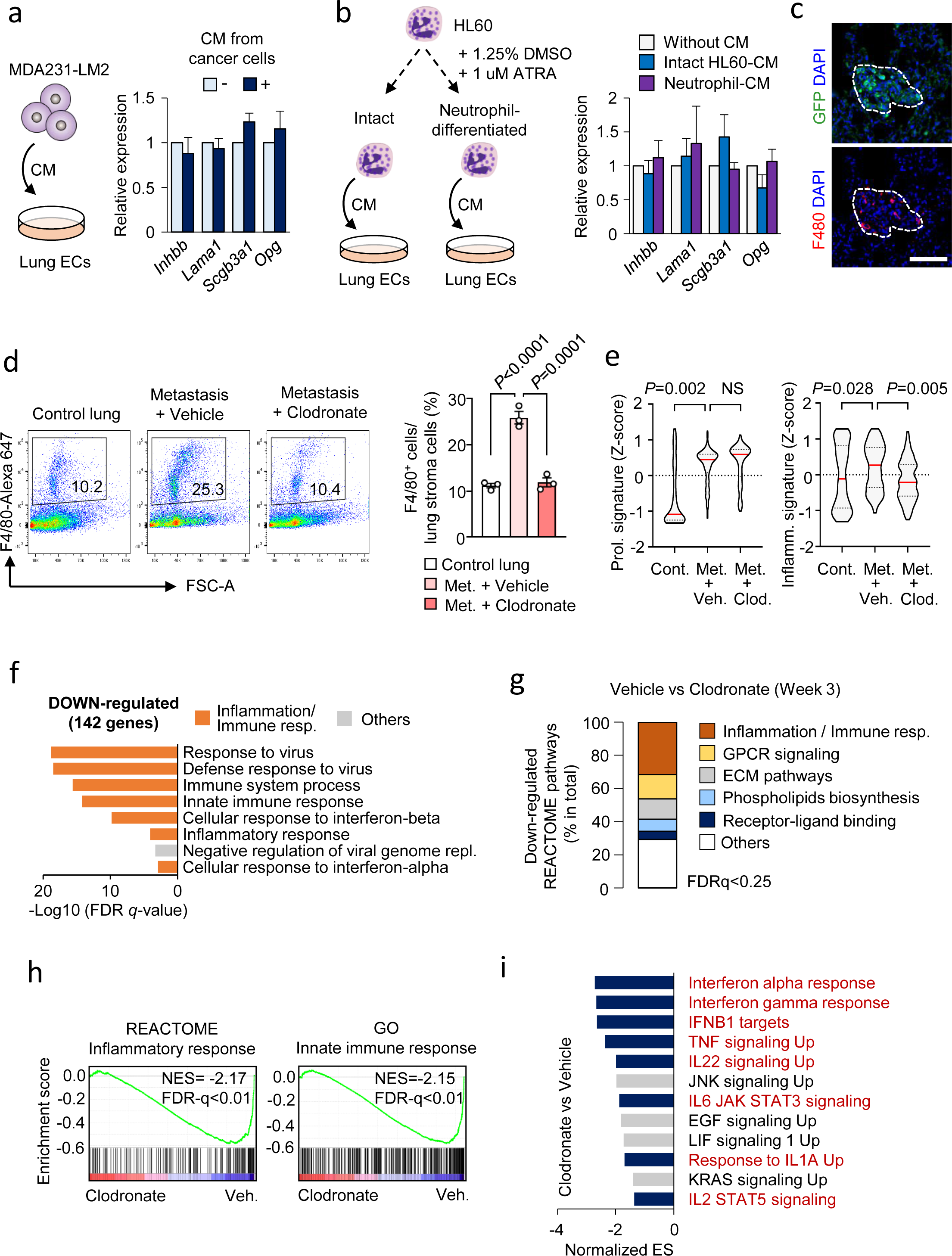
**a**, Expression of four vascular niche factors in human lung ECs treated with CM from MDA231-LM2 breast cancer cells. Schematic of the experiment (left) and relative expression (right). Values are mean from 3 or 4 independent experiments with SEM. **b**, Expression of the four vascular niche factors in human lung ECs treated with CM from intact HL60 cells or neutrophil-differentiated HL60 cells. Schematic setup of the experiment (left) and relative expression as means with SEM (right) from 3 or 4 independent experiments are shown. **c**, Immunofluorescence analysis of macrophages (F4/80) in metastatic foci in lung 2 weeks after the intravenous injection of MDA231-LM2 cancer cells (GFP). DAPI was used to stain nuclei. Scale bar, 100 μm. **d,** FACS analysis of F4/80^+^ macrophages in healthy control lung and week 3 metastatic lung treated with PBS-liposome or clodronate-liposome. Gating of FACS analysis (left) and quantified F4/80^+^ macrophages within lung stroma cells (right); *n* = 3 mice for each group. *P* values were determined by one-way ANOVA with Dunnet’s multiple comparison test. **e**, Z-scores of genes from a proliferation signature (Cell cycle mitotic, REACTOME) (left) and inflammation signature (Inflammatory response, Hallmark in MSigDB) (right) expressed in isolated ECs from metastatic lungs under the indicated conditions. *P* values were calculated with averaged Z-score of genes in signatures by one-way ANOVA with Fisher’s LSD test. *n* = 3 each group. NS, not significant. **f**, GO term analysis of down-regulated genes (142 genes, log2FC < -0.75, FDR < 0.25) in lung ECs after macrophage-depletion. **g**, Composition of down-regulated REACTOME pathway gene clusters with FDR < 0.25. **h**, GSEA plots of gene signature “Inflammatory response” (C2 in MSigDB, left) and “Innate immune response” (C5 in MSigDB, right) comparing lung ECs from mice treated with clodronate-liposome and PBS-liposome. NES, normalized enrichment score. **i**, Downregulated gene clusters of cellular signaling in lung ECs isolated from metastasis-bearing mice after macrophage depletion. GSEA was performed using Hallmark and C2 CPG in MSigDB. Signatures with FDR < 0.25 are shown. Inflammation-related signaling is highlighted with dark blue bars with red descriptions. ES, enrichment score.

**Supplementary Figure 7.**
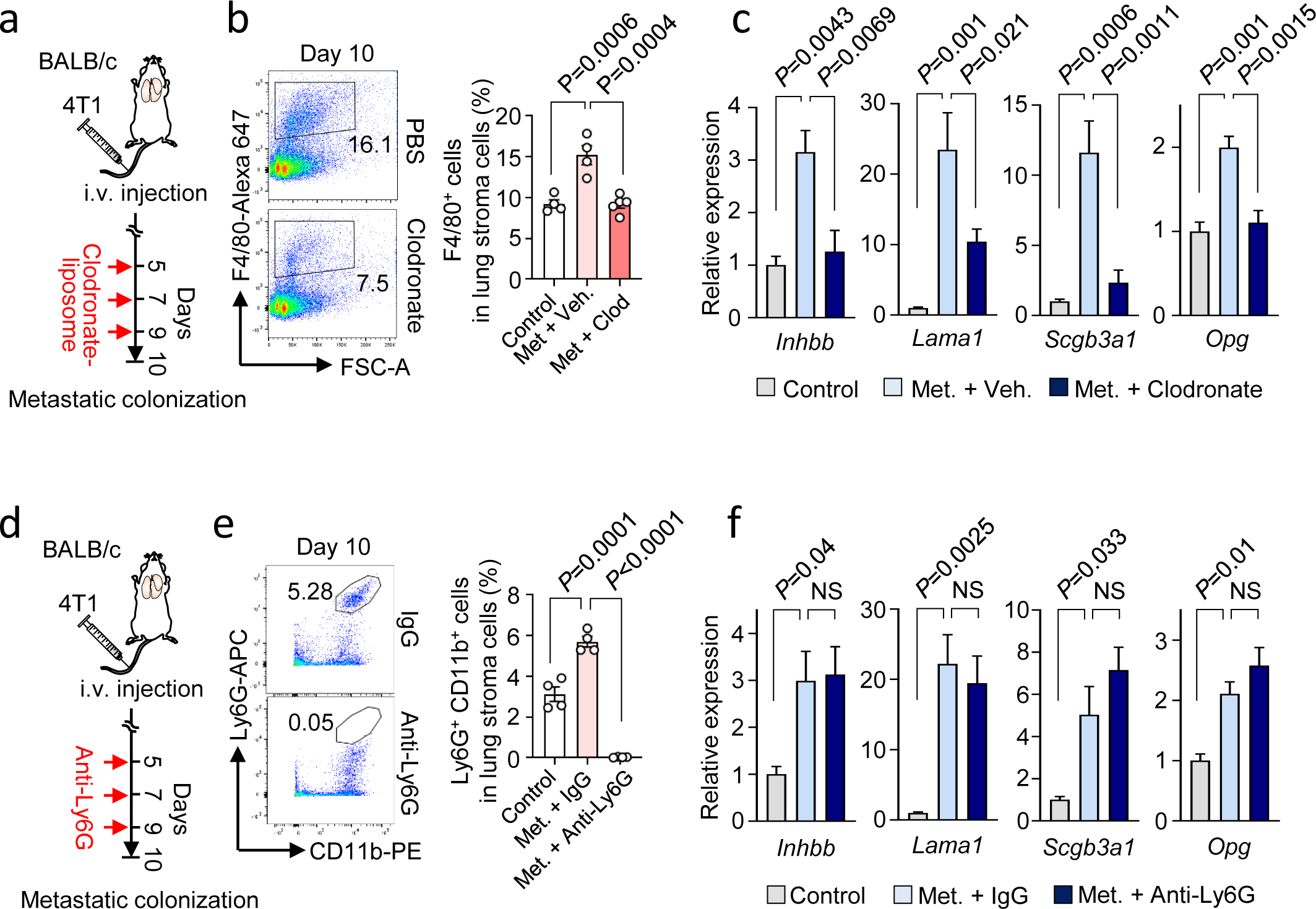
**a**, Scheme of macrophage-depletion by transient treatment with clodronate-liposome to BALB/c mice after intravenous injection of 4T1 cancer cells. **b**, FACS analysis of F4/80^+^ macrophages in metastatic lung treated with PBS-liposome or clodronate-liposome 10 days after the injection of cancer cells. *n* = 4 mice for each group. *P* values were determined by one-way ANOVA with Dunnet’s multiple comparison test. **c**, Expression of vascular niche components in lung ECs isolated from control healthy lung or metastatic lung (day 10) treated with PBS-liposome or clodronate-liposome; *n* = 4 or 5 mice. Mice harboring comparable lung metastatic loads, measured by *in vivo* bioluminescence imaging, were selected for analysis. *P* values were calculated by one-way ANOVA with Dunnet’s multiple comparison test. **d**, Scheme of the neutrophil-depletion by the transient treatment with anti-Ly6G antibody to BALB/c mice 10 days after the intravenous injection of 4T1 cells. **e**, FACS analysis of Ly6G^+^CD11b^+^ neutrophils in metastatic lung treated with IgG or anti-Ly6G antibody 10 days after the injection; *n* = 4 mice for each group. *P* values were determined by one-way ANOVA with Dunnet’s multiple comparison test. **f**, Expression of niche factors in lung ECs isolated from control healthy lung or metastatic lung (day 10) treated with IgG or anti-Ly6G antibody; *n* = 3 or 4 mice. Mice harboring comparable lung metastatic loads, measured by *in vivo* bioluminescence imaging, were selected for analysis. *P* values were calculated by one-way ANOVA with Dunnet’s multiple comparison test.

**Supplementary Figure 8.**
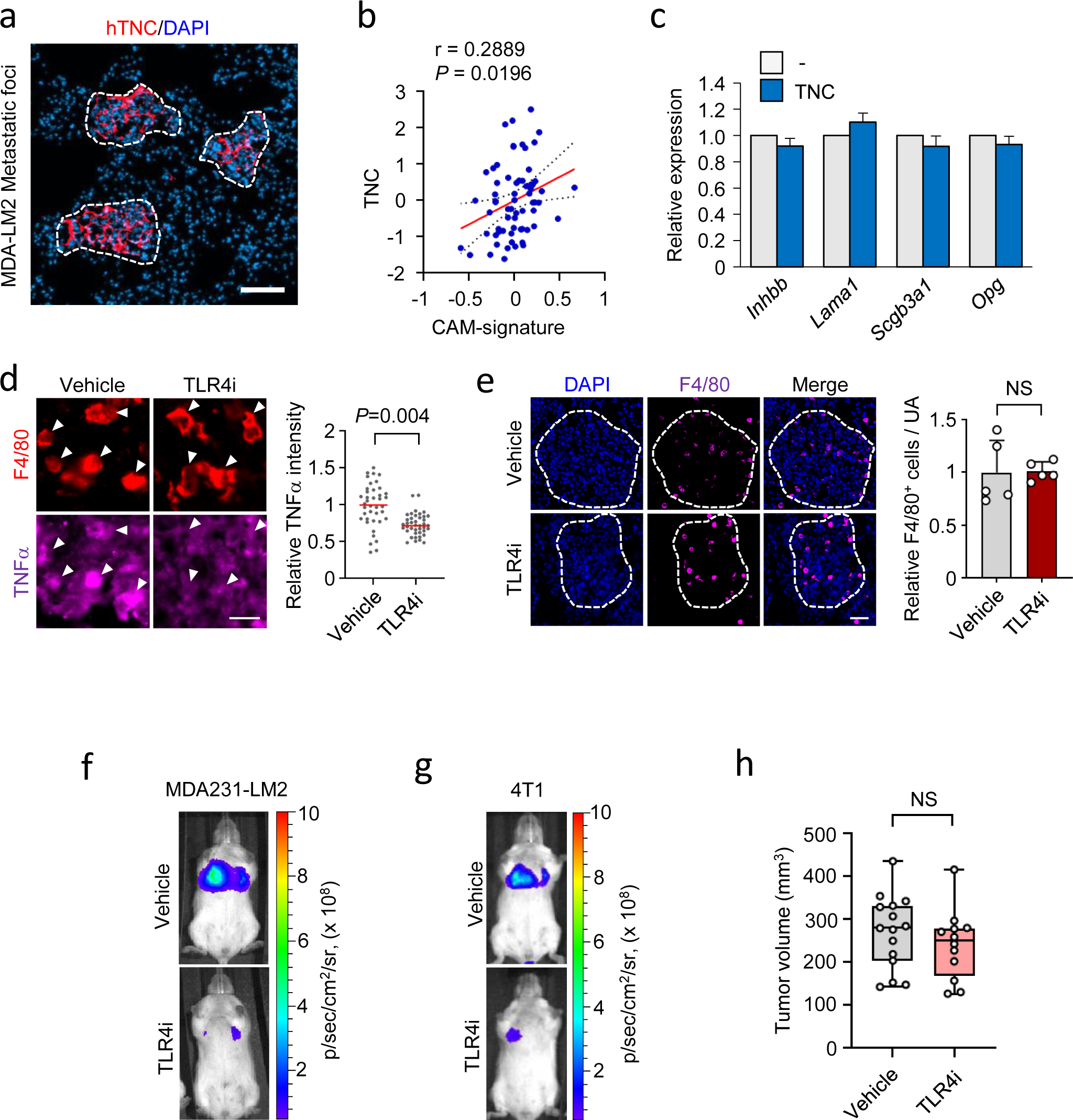
**a**, Immunofluorescence analysis showing accumulation of cancer cell-derived TNC in the metastatic foci at 2 weeks after intravenous injection of MDA231-LM2 cells. hTNC (red) and DAPI (blue). Scale bar, 100 μm. Dashed lines indicate the margins of metastatic foci. **b**, Correlation analysis of TNC and classically activated macrophage-signature (CAM-S) in 65 metastases samples from breast cancer patients. Linear regression with Pearson correlation r and two-tailed *P* values are shown. **c**, Expression of niche factors in lung ECs stimulated with 2 μg/ml of recombinant TNC for 20-24h *in vitro*. **d**, Immunofluorescence analysis of TNFα in F4/80^+^ macrophages within MDA231-LM2 metastatic nodules from mice treated with vehicle or TLR4i. Representative examples (left) and quantification of relative signals (right). Intensity of TNFα staining (purple) within F4/80 positive region (red) in the metastatic foci was measured; 40 and 41 macrophages, respectively, in lung from 5 mice were analyzed. *P* value was determined by one-tailed Mann-Whitney test. *n* = 5. Scale bar, 20 μm. **e**, Immunofluorescence analysis of macrophages (F4/80, purple) in MDA231-LM2 lung metastases from mice treated with vehicle or TLR4i. Dashed lines indicate the margins of metastatic foci. Relative number of F4/80^+^ cells in unit area of metastatic foci was quantified (right). Statistical analysis was performed with one-tailed Mann-Whitney test. *n* = 5. Scale bar, 50 μm. **f,g**, Representative examples of lung bioluminescence from Figure 7h. **h**, Volume of primary tumors measured on day 24 as in Figure 7j; vehicle: *n* = 15 tumors, TLR4i: *n* = 12 tumors. *P* values were calculated by one-tailed Mann-Whitney test.

**Supplementary Table 1.** GO term analysis of genes up-regulated in ECs associated with lung metastasis compared to control ECs in healthy lungs.

| Time point | Enriched GO term | P value | FDR | Fold enrichment |
| --- | --- | --- | --- | --- |
| Week1 | GO:0045087~innate immune response | 6.05E-07 | 5.83E-04 | 27.123 |
|  | GO:0009615~response to virus | 7.96E-06 | 0.0076 | 86.104 |
|  | GO:0002376~immune system process | 2.30E-05 | 0.0221 | 23.605 |
| Week2 | GO:0051607~defense response to virus | 1.28E-13 | 1.42E-10 | 64.965 |
|  | GO:0009615~response to virus | 2.44E-11 | 2.71E-08 | 100.455 |
|  | GO:0045087~innate immune response | 7.39E-09 | 8.18E-06 | 24.109 |
|  | GO:0002376~immune system process | 2.26E-07 | 2.59E-04 | 22.032 |
| Week3 | GO:0007049~cell cycle | 2.05E-37 | 3.31E-34 | 8.475 |
|  | GO:0051301~cell division | 1.46E-26 | 2.36E-23 | 9.433 |
|  | GO:0007067~mitotic nuclear division | 1.51E-25 | 2.44E-22 | 11.145 |
|  | GO:0051607~defense response to virus | 6.67E-12 | 1.08E-08 | 9.507 |
|  | GO:0006260~DNA replication | 8.76E-12 | 1.41E-08 | 11.470 |
|  | GO:0009615~response to virus | 1.01E-11 | 1.63E-08 | 14.700 |
|  | GO:0007059~chromosome segregation | 3.41E-10 | 5.50E-07 | 12.883 |
|  | GO:0002376~immune system process | 2.52E-09 | 4.07E-06 | 5.066 |
|  | GO:0045087~innate immune response | 5.42E-09 | 8.73E-06 | 4.851 |
|  | GO:0000281~mitotic cytokinesis | 3.43E-08 | 5.53E-05 | 23.521 |
|  | GO:0051310~metaphase plate congression | 1.29E-07 | 2.08E-04 | 44.102 |
|  | GO:0006270~DNA replication initiation | 2.18E-07 | 3.51E-04 | 25.726 |
|  | GO:0008283~cell proliferation | 1.34E-06 | 0.0021 | 5.613 |
|  | GO:0006268~DNA unwinding involved in DNA replication | 3.13E-06 | 0.0050 | 44.102 |
|  | GO:0000070~mitotic sister chromatid segregation | 4.96E-06 | 0.0080 | 23.009 |
|  | GO:0006974~cellular response to DNA damage stimulus | 5.92E-06 | 0.0095 | 3.780 |
|  | GO:0035458~cellular response to interferon-beta | 6.20E-06 | 0.0099 | 15.059 |
|  | GO:0045071~negative regulation of viral genome replication | 2.74E-05 | 0.0441 | 16.538 |

**Supplementary Table 2.**
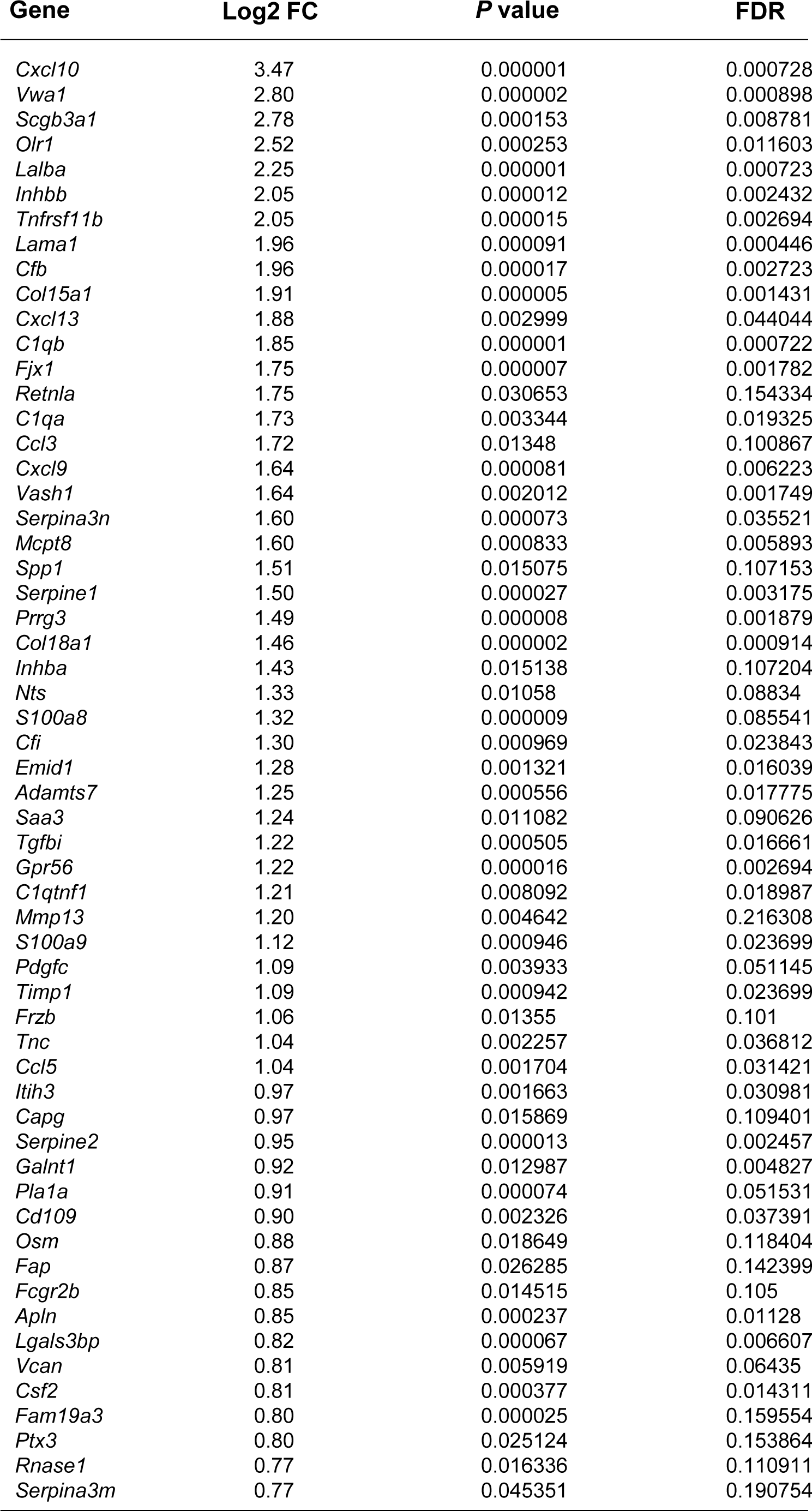
List of 58 genes of secreted proteins (GSP58) that are induced in metastasis-associating lung ECs at week 3. Genes with log2FC > 0.75, *P* < 0.05 and FDR < 0.25 are shown.

**Supplementary Table 3.**
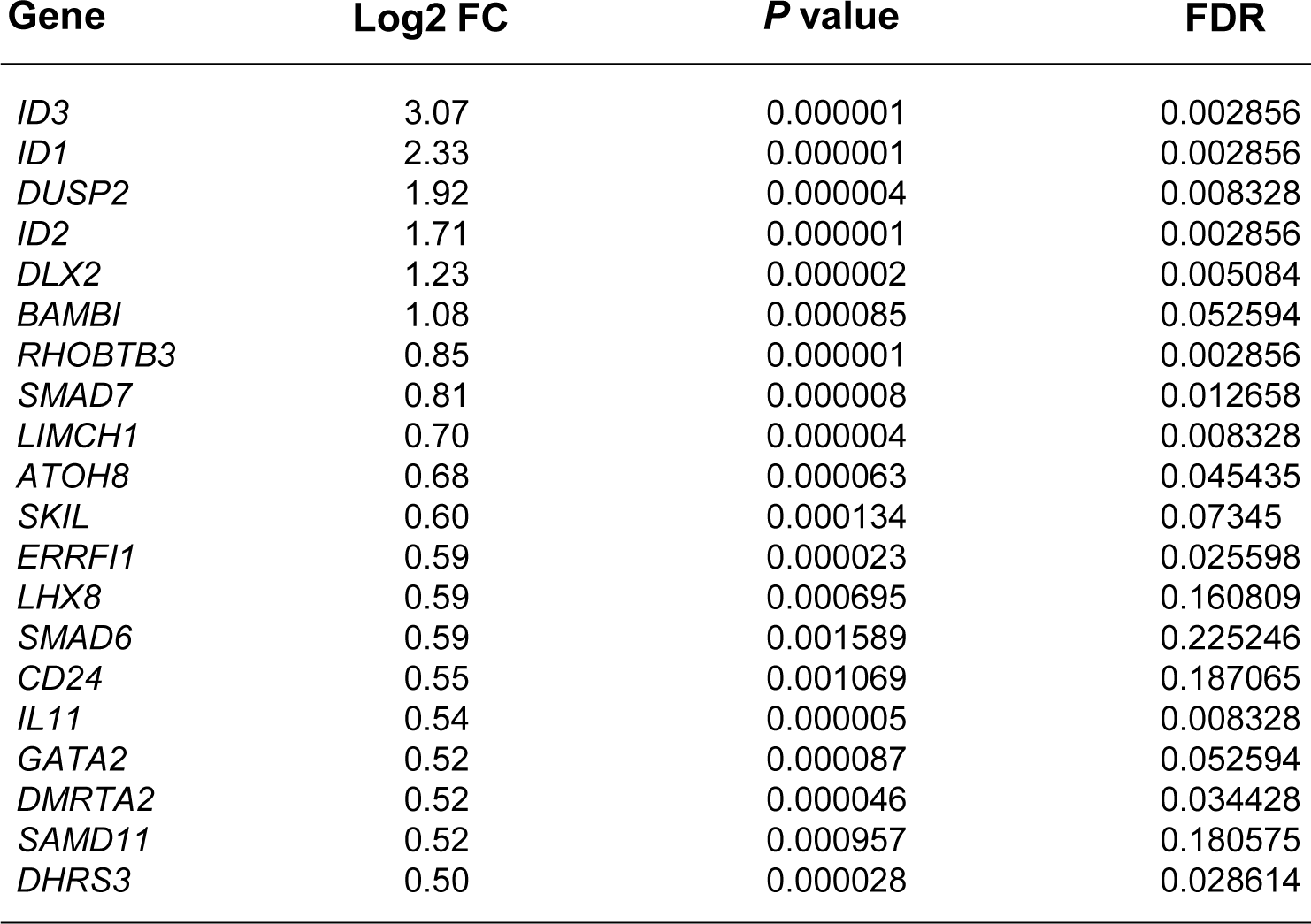
List of top 20 activin B-induced genes in breast cancer cells (activin B signature). Genes with log2FC > 0.5, *P* < 0.05 and FDR < 0.25 are shown.

| Gene | Log2 FC | P value | FDR |
| --- | --- | --- | --- |
| *ID3* | 3.07 | 0.000001 | 0.002856 |
| *ID1* | 2.33 | 0.000001 | 0.002856 |
| *DUSP2* | 1.92 | 0.000004 | 0.008328 |
| *ID2* | 1.71 | 0.000001 | 0.002856 |
| *DLX2* | 1.23 | 0.000002 | 0.005084 |
| *BAMBI* | 1.08 | 0.000085 | 0.052594 |
| *RHOBTB3* | 0.85 | 0.000001 | 0.002856 |
| *SMAD7* | 0.81 | 0.000008 | 0.012658 |
| *LIMCH1* | 0.70 | 0.000004 | 0.008328 |
| *ATOH8* | 0.68 | 0.000063 | 0.045435 |
| *SKIL* | 0.60 | 0.000134 | 0.07345 |
| *ERRFI1* | 0.59 | 0.000023 | 0.025598 |
| *LHX8* | 0.59 | 0.000695 | 0.160809 |
| *SMAD6* | 0.59 | 0.001589 | 0.225246 |
| *CD24* | 0.55 | 0.001069 | 0.187065 |
| *IL11* | 0.54 | 0.000005 | 0.008328 |
| *GATA2* | 0.52 | 0.000087 | 0.052594 |
| *DMRTA2* | 0.52 | 0.000046 | 0.034428 |
| *SAMD11* | 0.52 | 0.000957 | 0.180575 |
| *DHRS3* | 0.50 | 0.000028 | 0.028614 |

**Supplementary Table 4.**
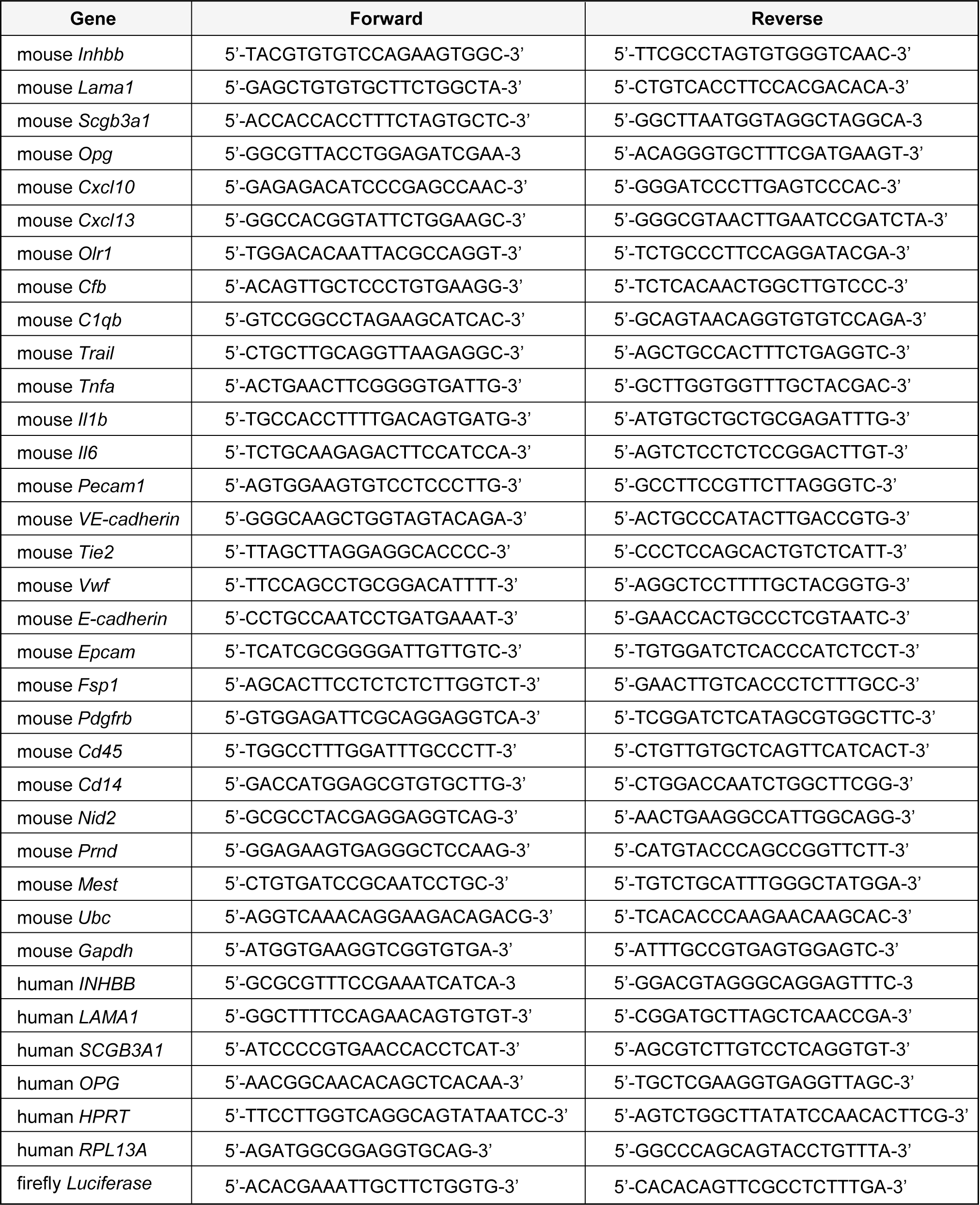
Primers used for qPCR.

## Notes

### Competing Interest Statement

The authors have declared no competing interest.

